# Cells dynamically adapt their nuclear volumes and proliferation rates during single to multicellular transitions

**DOI:** 10.1101/2025.10.04.679984

**Authors:** Vaibhav Mahajan, Keshav Gajendra Babu, Markus Mukenhirn, Antje Garside, Vinita Ajit Kini, Trishla Adhikari, Timon Beck, Byung Ho Lee, Kyoohyun Kim, Carsten Werner, Alf Honigmann, Sebastian Aland, Raimund Schlüßler, Anna Verena Taubenberger

## Abstract

Tumour development and progression are associated with biophysical alterations that manifest across multiple spatial scales, from the subcellular to multicellular tissue scale. While cells dynamically regulate their biophysical properties like volumes and mechanics in dependence of cell state and function, it is unclear how these properties are controlled in the dense multicellular environment of a developing tumour. Here, we quantified cell and nuclear volumes of single cancer cells, while they grew into multicellular tumour spheroids within well-defined, tuneable biohybrid polymer hydrogels. We quantitatively showed that the formation of multicellular structures is associated with marked reductions of cellular and nuclear volumes, cell cycle delays as well as cell mechanical alterations, and that these changes are coupled. Single-to-multicellular transitions coincided with a drastic decrease in median nuclear volumes by up to 60%, as well as overall cell volume decrease. Nuclear volume decrease could not be explained by compression due to confining microenvironments. Instead, cell cycle adaptions were one significant contributor, with smaller-sized G1 cells accumulating in growing clusters, an effect that was reversed by CDK1 inhibition. Another contributor was nuclear volume decrease in cells within clusters that was associated with higher mass density and stiffness and could be abrogated upon cell release from clusters. In turn, multicellular-to-single cell transitions that happened in cells that invaded from a tumour spheroid into the surrounding matrix, were accompanied by nuclear volume increases and cell softening. Taken together, our study provides insights into how cells dynamically adapt their cellular/nuclear volumes, cell cycle progression and mechanics in dependence of the multicellular state.

## Introduction

Cell volumes vary considerably across different tissues and cell types, as well as cell states and cellular processes. In early embryos, during cleavage stage, a multicellular organism forms from a single cell through a series of mitotic divisions, giving rise to smaller and smaller sized cells^1^. Cell differentiation can also be associated with drastic changes in cell volume^2–4^, as in the case of adipocytes whose volume can increase up to 30-fold compared to their precursors^5^. Moreover, cellular senescence and disease states like cancer are linked to critical changes in cell volume, underpinning its relevance in cellular function^6,7^. Given these and many more examples, it is essential for the cell to dynamically adapt its volume to a certain state that may be determined by a specific cellular function or by extracellular factors that it is exposed to within a particular microenvironment^8^.

In actively dividing cells, the cell volume naturally fluctuates with progression through the cell cycle, due to coordinated cell growth and division^9,10^. Before cell division, the cell increases its mass and volume while replicating DNA and synthesizing new cellular material^11,10^-subsequent division reduces cell volumes to about half^12^. Yet, cells are typically able to maintain a homogenous size distribution across multiple cell generations, which has been the subject of many studies. Mammalian cells were shown to exhibit so called ‘adder-like’ behaviour, where cells increase their volume by a defined amount over the cell cycle, in contrast to ‘sizer-like’ cells that increase their volume until a certain size threshold before dividing^12–14^. For effective volume adaption, the duration of the cell cycle phases^12^ as well as the growth rate itself can be tuned, e.g. via modulating biosynthesis rates^15,16^. To be able to adjust their growth kinetics, cells need to gauge their size and send feedback to the molecular machinery affecting volume change^10,12,17,18^. Such a mechanism can rely on volume-dependent dilution or concentration of molecular factors that positively or negatively regulate cell cycle progression, e.g. as shown for retinoblastoma protein^17^, cyclins and cyclin dependent kinases (CDKs)^17–19^. Cell growth rates along the cell cycle are also modulated by external factors, e.g. abundance of nutrients and growth factors^20^. For example, mTORC signalling regulates mRNA translation and *de novo* lipid synthesis and thereby can control cell growth in coordination with environmental conditions^21^.

Alterations in cell volume can also be linked to physical forces. For instance, cell volumes can change due to water flux that is driven by hydrostatic and osmotic pressure differences and is enabled by membrane permeability and facilitated diffusion through aquaporins^22^. By actively transporting ions or small osmolytes and through regulation of cortical tension, cells can control their intracellular pressure and volume^22–24^ and also drive cell shape changes, e.g. during mitosis^25^. Vice versa, shape changes, as occurring during cell spreading on a surface or when cells are under physical confinement^26,27^, can lead to changes in cell volume^28,29^. These force-induced changes in volume can also guide changes in cell proliferation and influence differentiation programs^30^. Experiments exposing cells to hyper- and hypoosmotic environments show that cells can adapt their volumes at short timescales (seconds to minutes), e.g. by regulatory volume increase or decrease, respectively^11,29,31–33^. Recently, mechanistic insights into volume regulation and coupled changes in membrane tension upon osmotic changes were shown, involving mTOR signalling, cytoskeleton control, and mechanosensitive ion transporters and channels^34^. Still, there is not much known about how cells regulate their volumes on a longer timescale when forming more complex tissue-like structures and when exposed to changing mechanical cues within 3D environments.

Size homeostasis is evidently perturbed in cancer cells. Within tumour tissues, cells are often found to be heterogeneous and overly enlarged in size^8^. This can partly be explained by polyploidy, which is known to affect cell volume^35^. Also, above-mentioned regulatory machinery of cell growth and proliferation control is commonly corrupted in cancer^36^. In addition, the tumour microenvironment represents a physically aberrant environment with stiff matrix, high cellular densities and the emergence of solid stress. As recently suggested, volume changes like swelling of cells, might be relevant during invasion^37,38^. Yet, it is still unclear how tumour cells regulate their volume as they proliferate to give rise to a multicellular structure in the 3D tissue context.

Here we grow multicellular structures from single cancer cells in a physiologically relevant, well-defined 3D hydrogel matrix to mimic early tumour formation. Using high resolution confocal and light sheet imaging combined with image segmentation, we show that single cells drastically reduce their cellular and nuclear volumes during formation of multicellular structures. The seen nuclear volume decreases could be partly attributed to changes in the cell cycle. As single cells grew into multicellular structures, cell cycle progression was delayed, leading to accumulation of intrinsically smaller G1 cells. Independent from cell cycle effects, however, cells within multicellular structures had increased mass density and higher Brillouin frequency shifts, indicating a more crowded and less compressible cytoplasm. Analogously, cell aggregates in suspension comprised of cells with lower nuclear volumes, suggesting that cell volume regulation differs between single cells and multicellular systems. Conversely, cells breaking away from multicellular structures at the onset of invasion adopted increased nuclear volumes again, suggesting that the process of volume adaption in dependence of the multicellular state is plastic. Overall, this study provides new insights into the regulation of cell volume in multicellularity and during processes relevant to tumorigenesis and invasion.

## Results

### Growth of tumour spheroids from single cells within defined 3D environments

We set out to study volume changes of single cells while they formed spheroids within extracellular matrix (ECM) mimicking hydrogels, as an *in-vitro* model of tumour microtissue formation **(Fig 1A)**. To that end, we embedded single MCF-7 breast cancer cells within mechanically tuneable MMP (matrix metalloprotease)-cleavable PEG-heparin hydrogels^39–41^ **(Fig 1A-D, Fig S1)**. To assess temporal morphological changes, we imaged cells after 1, 4 and 14 days in culture by confocal microscopy. While on day 1 single cells predominated, on day 4 small cell clusters of about 4-30 cells had formed **(Fig 1B)**. On day 14, larger sized spheroids with diameters of about 100-160 µm were observed, typically comprising more than 200 cells **(Fig 1C)**. Spheroids were densely packed with cells that were positive for ck8/18 and resembled the organization of mammary epithelial cells in clinical samples of ductal carcinomas *in-situ* (DCIS) **(Fig 1D)**.

**Figure 1.**
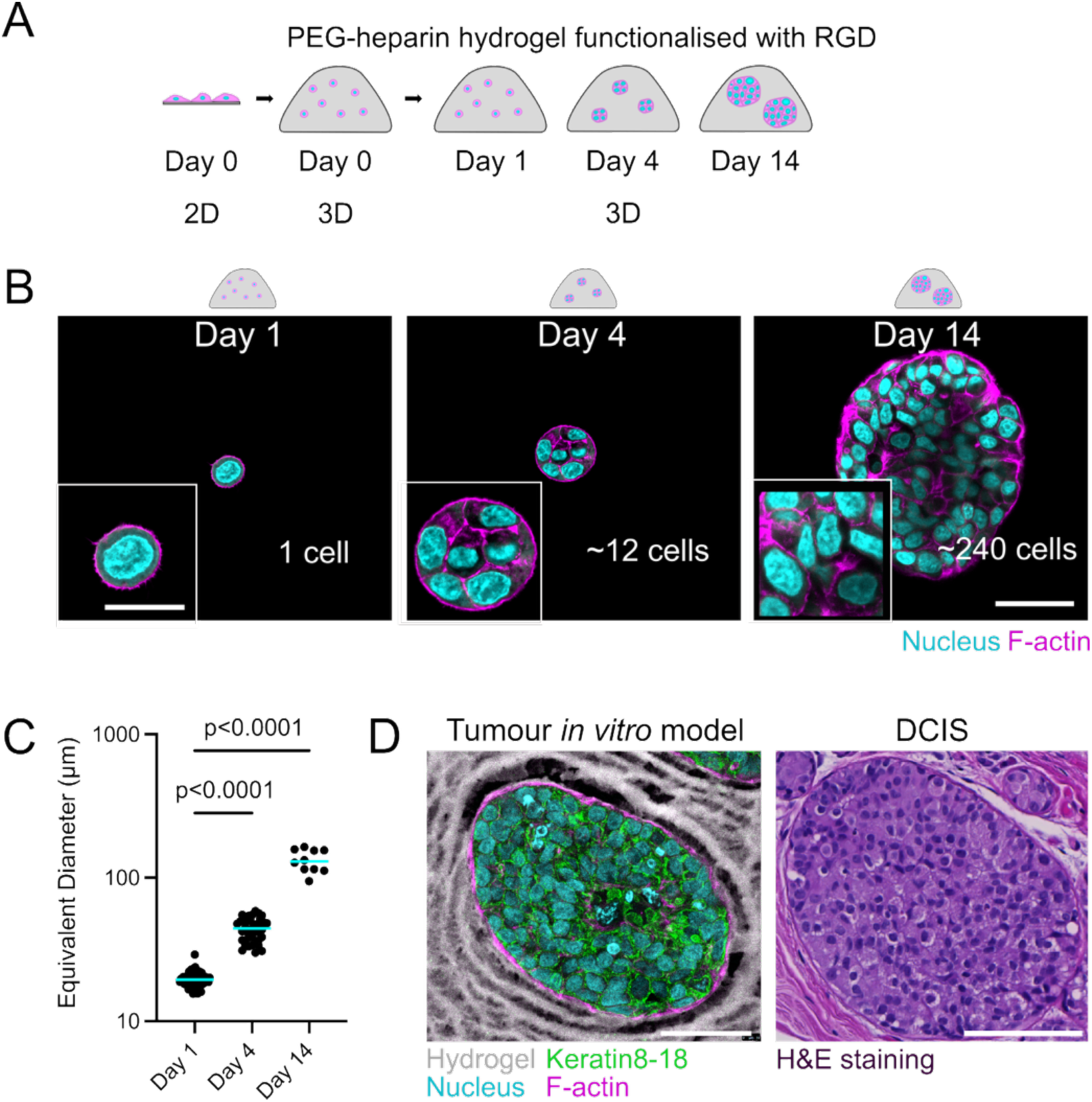
Forming multicellular structures from single cells within biohybrid polymer hydrogels. **(A)** Schematic illustrating the setup of the tumour microtissue in vitro model: Single cells are embedded into a compositionally and mechanically well-defined polymer hydrogel (PEG-heparin, elastic modulus 1.95 +/-0.15 kPa) and cultured as 3D culture for 14 days. **(B)** Confocal microscopy images of MCF-7 cells in compliant MMP-cleavable PEG-heparin gels stained with DAPI (nucleus) and Phalloidin-TRITC (F-actin) at day 1 (single cells), 4 (small clusters), 14 (spheroids). **(C)** Sizes of single cells (day 1), small clusters (day 4) and spheroids (day 14). An equivalent diameter was calculated from the object’s cross-sectional area assuming spherical object, n = 55 (day 1), 38 (day 4), 10 (day 14); 4-6 gels each; N = 2-3. Cyan bars represent the median. Kruskal-Wallis test with multiple comparisons (Dunn’s) was performed for statistical analysis. **(D)** Representative image of a MCF-7 tumour spheroid and its surrounding hydrogel matrix (labelled by fluorescent maleimide). Frozen sections were stained with DAPI (nucleus), Phalloidin-TRITC (F-actin), and keratin 8-18. Note that cryofreezing makes the gel macroporous^42^. (Right) H&E-stained section of a patient-derived ductal carcinoma in-situ (DCIS) tissue specimen. Scale bars – (B) 50 µm and 25 µm inset; (D) 50 µm (Left) and 100 µm (right).

### Nuclear volumes decrease in spheroids compared to single cells

Following establishment of the 3D culture model, we labelled the plasma membranes of live MCF-7 H2B-mCherry cells growing in 3D and performed spinning disk confocal microscopy at the above-chosen time points. **(Fig 2A)**. After 3D segmentation of nuclei and cell bodies/whole clusters from resolved z-stacks by StarDist and Limeseg, respectively, we calculated the cells’ nuclear and cellular volumes **(Fig 2B, C)**. Surprisingly, the median nuclear volume of MCF-7 H2B-mCherry cells decreased by ∼40% at day 4 and by ∼60% at day 14 compared to single cells at day 1 **(Fig 2D)**. Such pronounced reduction in nuclear volumes within spheroids versus single cells was also seen for PANC1 H2B-mCherry (pancreatic cancer) cells **(Fig 2E, Fig S2A)** and non-cancerous MCF10A H2B-GFP (breast epithelial) cells **(Fig 2F, Fig S2B)**, suggesting to us a more general phenomenon. We wondered whether the cell volumes were decreased within the multicellular clusters, too. While the volumes of single cells could be accurately quantified after membrane-based segmentation **(Fig 2B)**, this approach was not feasible for the denser spheroids due to the heterogenous cell shapes. Therefore, we rather determined the average cell volumes on day 4 and 14 by dividing whole cluster volumes by the constituent nuclei numbers. Concomitant with above-seen nuclear volume reductions, a clear decrease in the average cell volume was observed within MCF-7 H2B-mCherry spheroids (∼77%) **(Fig 2G)** and HeLa FUCCI cervical cancer small clusters (∼35%) **(Fig S2C)**, when compared to single cells. We further analysed the spatial distribution of nuclear volumes in MCF-7 H2B-mCherry spheroids (day 14) and found no systematic differences either in nuclear volumes or their nearest-neighbour-distances, when moving from spheroid core to periphery **(Fig S3)**.

**Figure 2.**
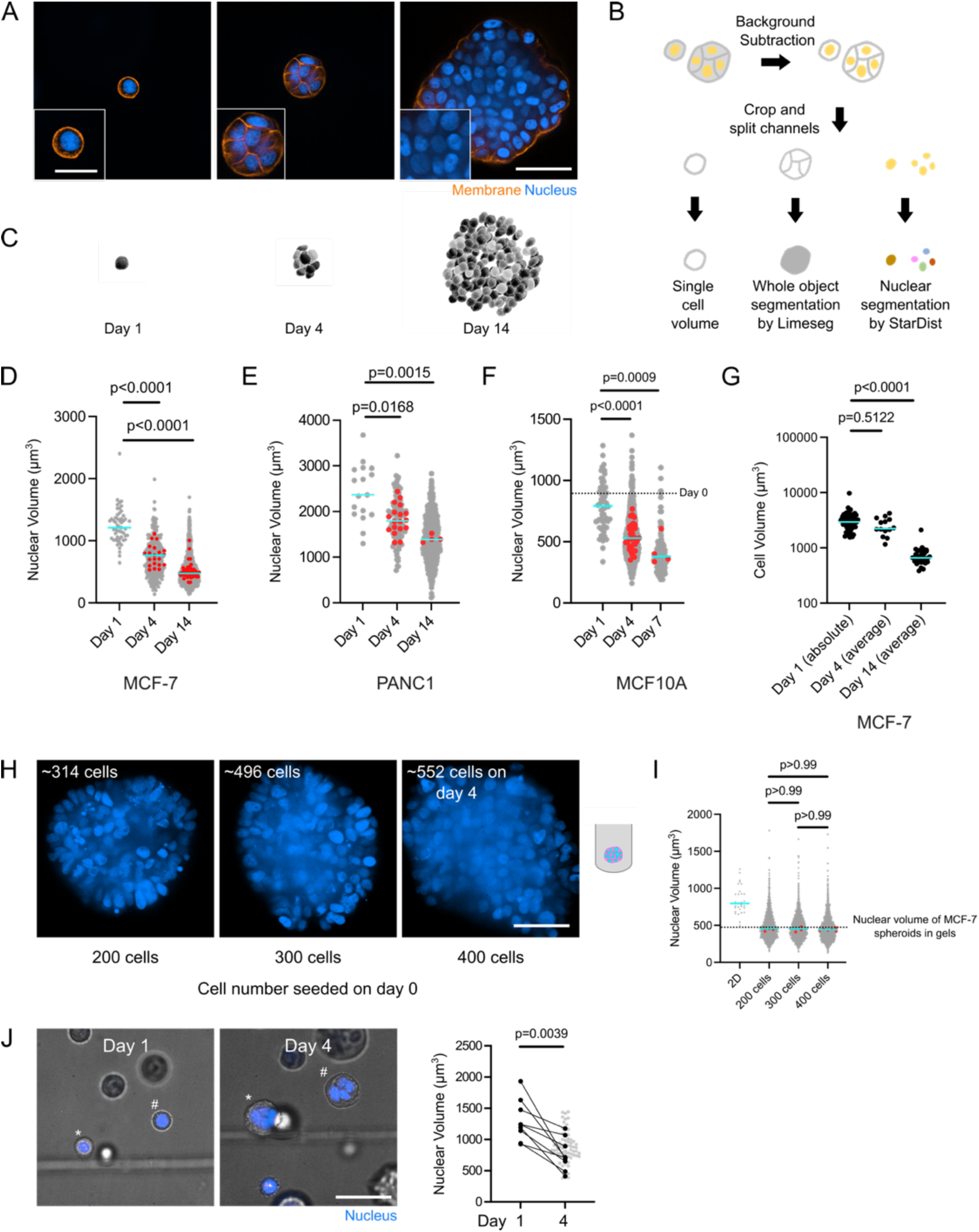
Cells forming multicellular structures show reductions in their nuclear and cellular volumes. **(A)** Live confocal images of MCF-7 H2B-mCherry in compliant MMP-cleavable PEG-heparin gels stained with CellMask Deep Red (membrane) at day 1, 4, 14. **(B)** Schematic showing image analysis methodology for quantifying cellular and nuclear volumes, as well as overall object volumes. **(C)** Nuclear segmentation of MCF-7 cells at day 1, 4, 14 from (A). **(D)** Scatter plot showing nuclear volumes of MCF-7 H2B-mCherry in gels at day 1, 4, 14. n = 53 cells (day 1), 141 cells from 21 clusters (day 4), 1609 cells from 10 spheroids represented as quarters (day 14); 4-6 gels each; N = 2-3. **(E)** Nuclear volumes of PANC1 H2B-mCherry in gels at day 1, 4, 14. n = 17 cells (day 1), 121 cells from 18 clusters (day 4), 695 cells from 4 spheroids (day 14); 4 gels each; N = 2. **(F)** Nuclear volumes of MCF10A H2B-GFP in gels at day 0, 1, 4, 7. n = average from 7 cells and represented as magenta dotted line (day 0), 60 cells from 39 doublets (day 1), 661 cells from 35 clusters (day 4), 142 cells from 4 spheroids (day 7); 2-4 gels each; N = 1-2. **(G)** Cell volumes for MCF-7 H2B-mCherry in gels at day 1, 4, 14. n = 55 (day 1), 14 (day 4), 32 (day 14); 4-6 gels each; N = 2-3. For single cells, whole cell could be segmented from the membrane signal. For multicellular structures, the whole cluster (for day 4) or quarter of a spheroid (for day 14) was segmented from the membrane signal and this was divided by the number of nuclei counted in each structure to obtain an average cell volume. **(H)** Live confocal images of MCF-7 H2B-mCherry as free-floating aggregates formed from 200, 300, 400 cells initially. At day 4 (upon aggregate formation), the final cell numbers were ∼314, ∼496, ∼552 cells, respectively. **(I)** Nuclear volumes of MCF-7 H2B-mCherry free-floating aggregates (from H) and for comparison resuspended single MCF-7 H2B-mCherry cells from 2D cultures. n = 31 cells (2D), 1590 cells (200 cells), 2426 cells (300 cells), 2477 cells (400 cells) from 6 aggregates each; N = 1. Magenta dotted line corresponds to median nuclear volume for MCF-7 H2B-mCherry spheroids in gels at day 14. **(J)** (Left) Live confocal images of the same MCF-7 H2B-mCherry cells imaged at day 1 and 4 in gels. (Right) Matched nuclear volume of MCF-7 H2B-mCherry in gels at day 1 and 4. n = 9; N = 1. For D-G, I: Cyan bars represent the median. Red dots correspond to the median value per structure. For D-G, I: Kruskal-Wallis test with multiple comparisons (Dunn’s) was performed for statistical analysis. For J: Wilcoxon matched-pairs signed rank test was performed for statistical analysis. Scale bars – (A) 50 µm and 25 µm inset; (H, J) 50 µm.

We next evaluated whether the seen nuclear volume changes also occurred in spheroids that had formed solely by cell aggregation. Thus, we measured nuclear volumes in free-floating spheroids that had assembled from differently sized cell numbers (200, 300 and 400 cells) over 4 days **(Fig 2H)** in low attachment u-shaped plates. Initial cell number appeared to have no effect on the nuclear volumes of the spheroids’ constituent cells. Compared to single resuspended cells from 2D cultures, however, the aggregates showed a ∼44% lower nuclear volume **(Fig 2I)**, which was equivalent to the nuclear volumes of day 14 spheroid cultures within the 3D matrix. Together, these results show that cells within multicellular structures have largely reduced cellular and nuclear volumes compared to single cells, independent of whether they are grown in a 3D matrix or as free-floating aggregates.

We noticed that the nuclear/cellular volume distributions only partially overlapped at subsequent timepoints **(Fig 2D-G, Fig S3C)**, so the smaller volumes were unlikely to arise from differently sized subpopulations of cells. Nevertheless, we compared nuclear volumes of MCF-7 H2B-mCherry single cells at day 1 and the corresponding cluster at day 4. Indeed, the nuclear volume of the initial cell was always larger compared to its day 4 progeny **(Fig 2J)**, so single cells evidently shrank in volume with culture time and/or formation of clusters.

### Nuclear volume reductions occur independently of mechanical confinement

Considering possible contributors to cell/nuclear volume decrease, we asked whether growth induced build-up of compressive stress and mechanical confinement of tumour spheroids may contribute to nuclear volume reductions. Thus, we decided to vary the level of compressive stress on forming MCF-7 H2B-mCherry spheroids (day 14) by tuning hydrogel stiffness and MMP-cleavability **(Fig S1, Fig 3A)** as previously shown (between 0 to 2.5 kPa)^40^. Although the spheroid sizes were sensitive to increased levels of compressive stress **(Fig S4)**, nuclear volumes of cells within spheroids were similar under all conditions **(Fig 3B)**, ruling out that cell compression within spheroids was responsible for the measured nuclear size changes. In addition, we interfered with mechanosensitive ion channels by treatment with GsMTx4, which also did not abolish nuclear volume decreases from day 1 to 4 **(Fig S5)**. Together, these results suggest that nuclear volume decreases occurred independently of cell confinement and mechanical stress sensed by cells within growing multicellular structures.

**Figure 3.**
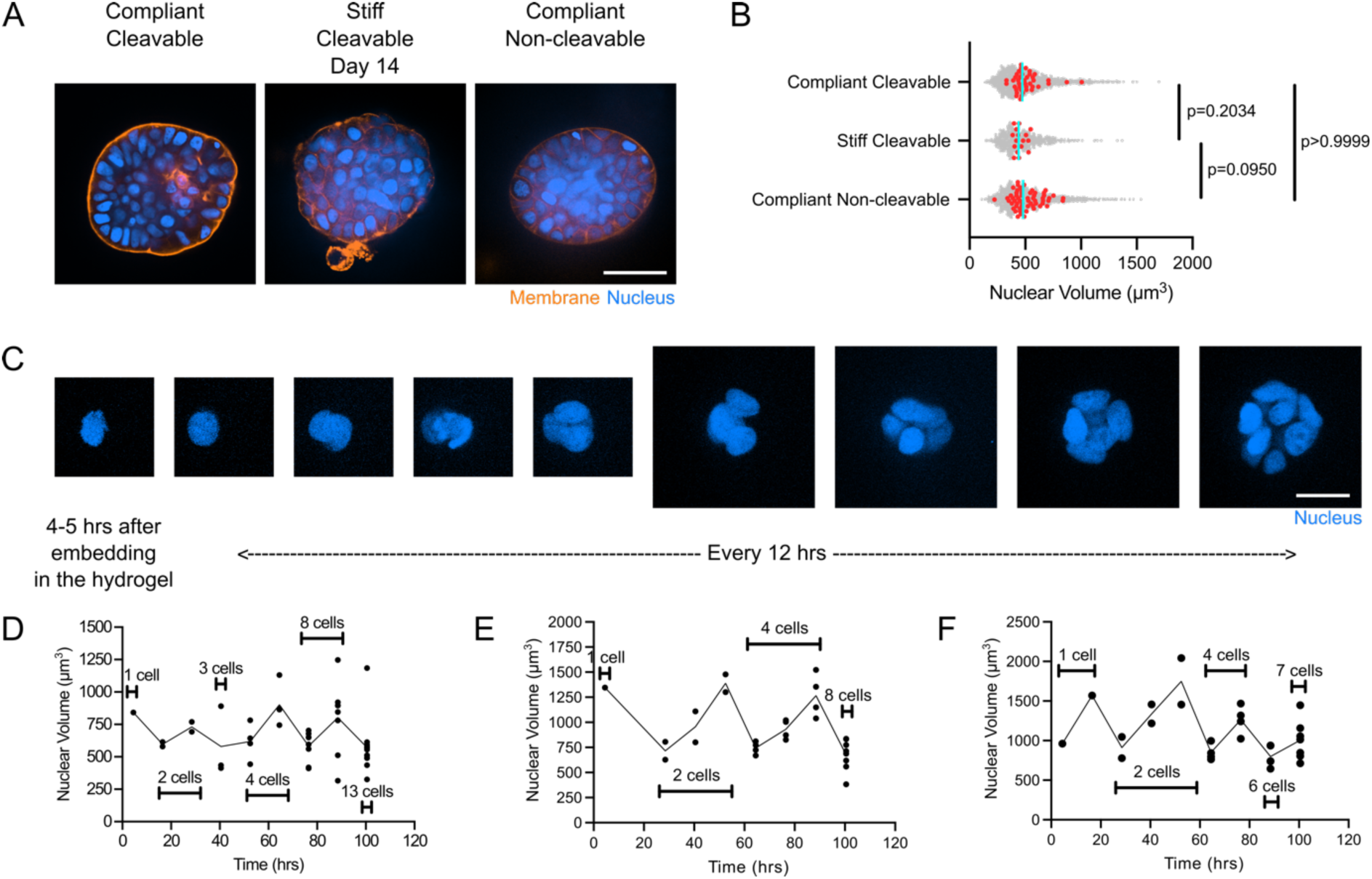
Nuclear volumes of multicellular structures are unaffected by compressive stress and alter over the cell cycle. **(A)** Live confocal images of MCF-7 H2B-mCherry at day 14 in compliant (1.95 +/- 0.15 kPa) and stiff (18.75 +/-2.18 kPa) MMP-cleavable and compliant non-cleavable (2.64 +/-0.29 kPa) PEG-heparin gels stained with CellMask Deep Red (membrane). **(B)** Scatter plots showing nuclear volumes of MCF-7 H2B-mCherry cells at day 14 in different PEG-heparin gel environments. N = 1609 cells from 10 spheroids represented as quarters (compliant MMP-cleavable), 709 cells from 6 spheroids represented as quarters (stiff cleavable), 344 cells from 15 spheroids represented as quarters (compliant non-cleavable); 4-6 gels each; N = 2-3. Cyan bars represent the median. Red dots correspond to the median value for one quarter of a spheroid. Kruskal-Wallis test with multiple comparisons (Dunn’s) was performed for statistical analysis. **(C)** (Left) Timelapse confocal microscopy (12-hour intervals) of MCF-7 H2B-mCherry in a gel starting from day 0 to 4. (Right) Nuclear volumes of MCF-7 H2B-mCherry from day 0 to 4 along with cell numbers at each timepoint corresponding to adjoining timelapse data. **(C-D)** Additional examples of nuclear volumes of MCF-7 H2B-mCherry in gels from day 0 to 4 at 12-hour intervals. Scale bars – (A) 50 µm; (C) 20 µm.

### The proportion of G1 cells increases in spheroids compared to single cells

Recently, cell volume reductions in dense cellular monolayers were reported due to size-reducing cell divisions upon contact inhibition^43^. This led us to monitor nuclear volumes longitudinally over time rather than only analysing temporal snapshots. As expected, we measured significant fluctuations of nuclear volumes over time along with cell cycle progression **(Fig 3C-E)**. Since the nuclear volume reductions from single cells (day 1) to small clusters (day 4) were within the rough range of volume fluctuations over the cell cycle, we decided to focus next on the dynamics of nuclear volume changes across the cell cycle. Therefore, we switched to cell lines expressing the FUCCI cell cycle reporter^44^. This allowed distinction of cell cycle stages based on alternating levels of orange/red [Cdt1(30/12)] fluorescent signal accumulating in G1 phase and green [Geminin(1/110)] signal in S/G2/M phases^45^ **(Fig 4A)**. While on day 1 most single MCF-7 FUCCI cells (∼86%) were in the S/G2 phase, on day 4 we observed ∼46% cells in G1 cell cycle phase **(Fig 4B)**. Mitotic cells were not counted, since they were not detected during segmentation due to absence of the nucleus. Quantitative analysis of nuclear volumes of MCF-FUCCI cells revealed significantly decreased nuclear volumes of G1 compared to S/G2 cells **(Fig 4C).** The same trend of increasing proportions of G1 cells within clusters was observed for HeLa cells **(Fig S6A)**, for MCF-7 FUCCI single cells when they were released from the gels at different timepoints **(Fig S6B)** and for MCF-7 FUCCI cells cultured as free-floating aggregates **(Fig S7)**. Concomitantly, bulk RNA sequencing data indicated changes in cell cycle related gene sets and genes between multicellular structures and single cells **(Fig S8)**.

**Figure 4:**
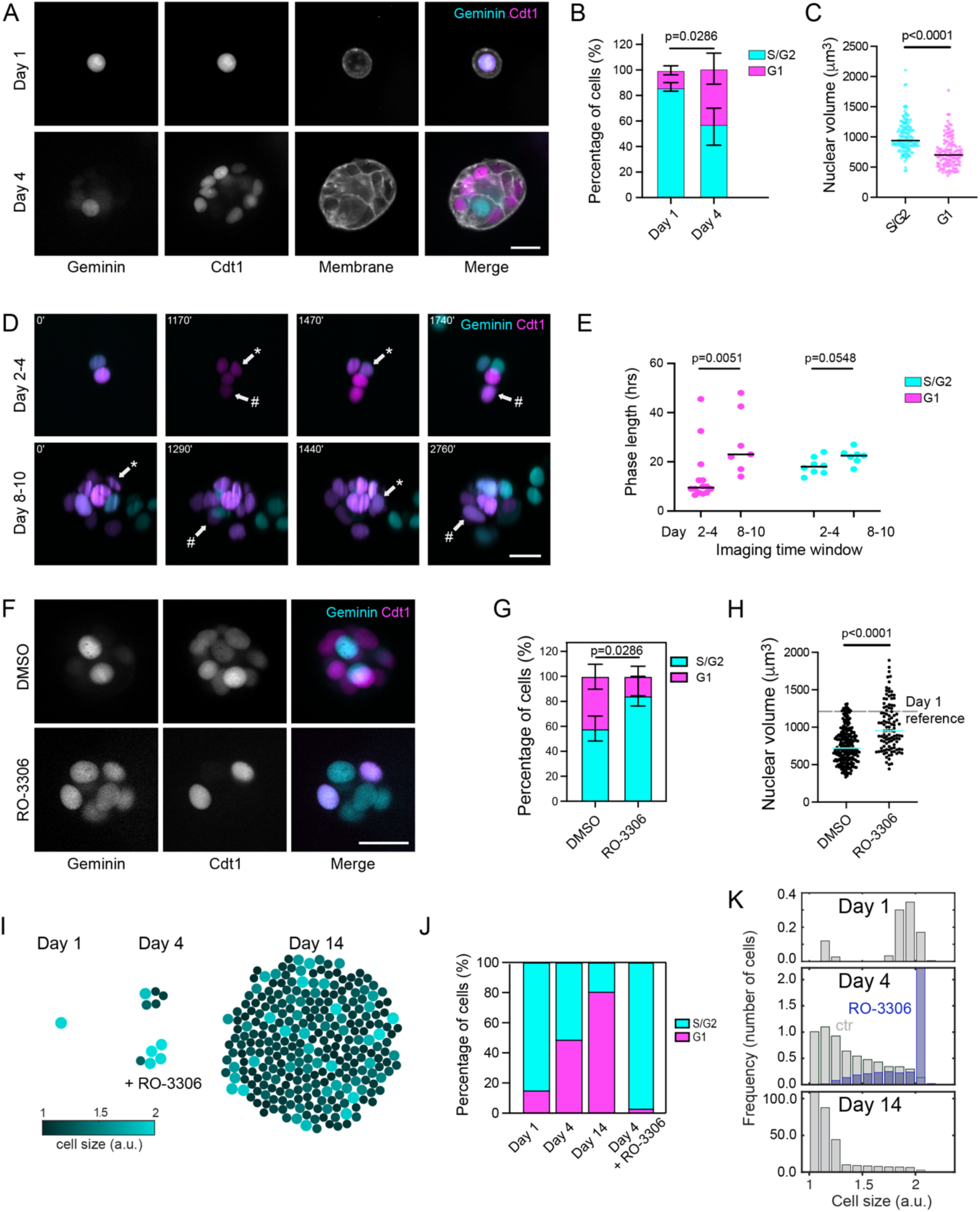
Accumulation of smaller G1 cells can partly account for the decrease in nuclear volume in multicellular structures. **(A)** Live confocal images of MCF-7 FUCCI cells in compliant MMP-cleavable PEG-heparin gels stained with CellMask Deep Red (membrane) at day 1 and 4. **(B)** Percentage of MCF-7 FUCCI cells in G1 and S/G2 phases in at day 1 and 4. n = 43 (day 1), 305 (day 4); 4 gels each; N = 2. Medians with range are shown. **(C)** Nuclear volumes of G1 and S/G2 MCF-7 FUCCI cells in small clusters (day 4) in gels. Black bars represent the median. n = 149 (S/G2), 125 (G1); 4 gels; N = 2. **(D)** Live fluorescent images of MCF-7 FUCCI cells imaged between day 2 to 4 and day 8 to 10 using light sheet microscopy. Arrows show start of G1 phase (Cdt1 only) and end of G1 phase (Geminin onset) for the same nuclei along with time stamps. **(E)** Cell cycle phase lengths from light sheet timelapse microscopy of MCF-7 FUCCI cells between days 2 to 4 and 8 to 10 of culture in compliant MMP-cleavable PEG-heparin gels. n = 14 (G1; day 2 to 4), 8 (G1; day 8 to 10), 7 (S/G2; day 2 to 4), 7 (S/G2; day 8 to 10); 2-4 entities each; N = 1. Black bars represent medians. **(F)** Live confocal images of MCF-7 FUCCI cells in gels after 24-hour treatment with 5 µm RO-3306 at day 4. **(G)** Percentage of MCF-7 FUCCI cells in G1 and S/G2 phases in gels at day 4 after treatment with RO-3306. n = 295 (DMSO), 184 (RO-3306); 4 gels; N = 2. Medians with range are shown. **(H)** Nuclear volumes of MCF-7 FUCCI cells within small clusters treated with RO-3306 at day 4 in gels. n = 238 (DMSO), 109 (RO-3306); 4 gels; N = 2. Cyan bars represent the median. **(I)** Numerical simulation of growing and dividing cells over time. Representative simulated cells and cell colony at day 1, 4, and 14 (colour coded for cell volume) are shown. **(J, K)** Cell cycle distribution within G1 and S/G2 phases (**J**) and volume (**K**) distribution of the resultant cell population at each time point (averaged over 1000 simulations). The action of RO-3306 is simulated after day 3 by stopping division and allowing growth and accumulation of cells in G2. For B, C, G, H: Mann-Whitney test was performed for statistical analysis. For E: Kruskal-Wallis test with multiple comparisons (Dunn’s) was performed for statistical analysis. Scale bars – (A, D, F) 25 µm.

### Changes in cell cycle distribution partly explain the decrease in nuclear volume

The seen accumulation of cells in G1 phase could either be due to prolongation of the G1 phase or the volume growth rate over G1 might be reduced over time. To better understand this, light sheet timelapse microscopy was performed on MCF-7 FUCCI cells from day 2 to 4 and then from day 8 to 10 **(Fig 4D)**. When comparing nuclear orange/green fluorescent signals in segmented z-stacks or over z-projections, we estimated a median G1 length of ∼10 hours for cells imaged between day 2 and 4. In contrast, at the later time point between day 8 and 10, median G1 lengths were significantly prolonged to at least 23 hours **(Fig 4E**, **Table 1**). At the same time, S/G2 phase lengths did not show any significant changes, though it increased from ∼18 hours (day 2 to 4) to at least 23 hours (day 8 to 10) **(Fig 4E**, **Table 1**). Still, all cell cycle stages were seen within larger clusters indicating that cells progressed through the cell cycle in principle. Thus, we concluded that during later time points in culture, cell cycle progression is slowed down primarily due to lengthening of G1 phase but does not stall.

We wondered whether the higher percentage of naturally smaller G1 cell nuclei compared to S/G2 cells **(Fig 4C)** may be sufficient to explain the overall nuclear volume reductions in multicellular structures. To investigate how shifts in cell cycle distribution affected nuclear volumes, we treated MCF-7 FUCCI clusters on day 3 with RO-3306, a CDK1 inhibitor that arrests cells in S/G2 phase^46^. As expected, we noted an accumulation of S/G2 cells by day 4, after 24 hrs of treatment **(Fig 4F, G)**. This shift in cell cycle distribution was associated with a partial recovery from the above-described nuclear volume decrease **(Fig 4H)**, suggesting that the seen changes in cell cycle distribution contributed to the decreasing nuclear volumes during spheroid formation.

To support these findings, we built a simple theoretical model capturing cell-cycle related volume changes of a population of initially single cells proliferating into spheroids. The model was fed with growth rates that were experimentally measured at an early timepoint (between days 1 and 4) using light-sheet timelapse microscopy. At the start of each cell cycle, cells were modelled to increase their volumes exponentially until reaching a size threshold of double their initial cell size, then dividing into half. In the model, we further considered the above seen lengthening of G1 cell cycle phase with cluster size, while G2 remained constant. The model emulates the experimental findings of an increasing proportion of smaller sized cells that progressively lower the average nuclear volume of the cell population **(Fig 4H)**. We further simulated the effect of RO-3306 in our numerical simulations, starting from day 3 on, where cell cycle progression was halted at the end of G2, but cells continued to increase in volume. As expected, this resulted in an increased proportion of S/G2 cells at day 4 and a concomitant increase of the average cell volume **(Fig 4I)**.

Thus, both the experiments and theoretical model show that a progressive cell cycle shift can account for the seen cellular and nuclear volume decreases over culture time. Nevertheless, opposed to the theoretical model, experimental accumulation of S/G2 cells did not completely restore the reduction in nuclear volume compared to single cells (day 1) **(Fig 4G, I)**, hinting at an additional factor causing nuclear volume changes with multicellularity.

### Cell release from spheroids results in cell swelling

We noted a discrepancy in nuclear volumes between cells that had been released from multicellular structures and their *in-situ* measured volumes **(Fig S6B)**. Thus, we systematically compared nuclear volumes of MCF-7 H2B-mCherry cells before and after release and dispersion into single cells at day 1, 4 and 14 **(Fig 5A)**. While nuclear volumes were similar at day 1 when measured *in-situ* (i.e. within hydrogel) and *ex-situ* (in suspension after release) **(Fig 5B)**, median nuclear volumes of released cells were significantly increased by ∼46% and ∼95% when compared to their *in-situ* counterparts at day 4 and 14, respectively **(Fig 5B)**. Since the release and imaging process was completed within less than one hour, we excluded nuclear volume increases due to further cell cycle progression during the release and imaging period as a possible reason. Despite the seen nuclear volume recovery upon release, significant differences in the median nuclear and cellular volumes remained for released cells over culture time **(Fig 4B, Fig S9)**. We thought that this was likely to be attributed to above-described cell cycle shifts. However, when directly comparing cells of the same cell cycle stage, we still noted a minor reduction in nuclear volume of released S/G2 cells at day 14 (∼20%) compared to day 1, which might be attributed to different volumes from early to late G2 stage or due to an incomplete recovery process **(Fig 5C)**.

**Figure 5:**
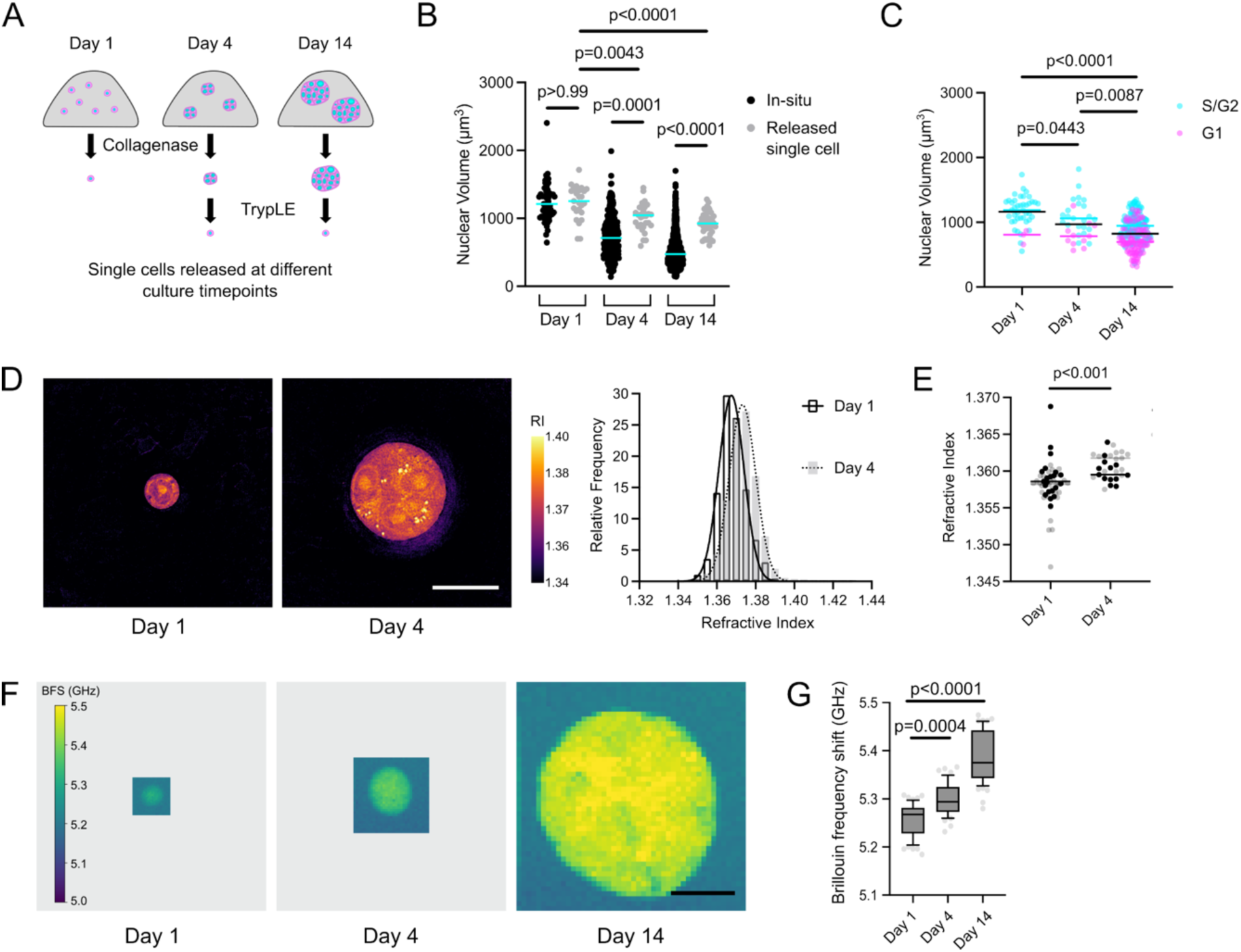
Cells within multicellular structures have a more crowded cytoplasm and decreased compressibility. **(A)** Schematic showing release of single cells (from gel and formed clusters) at different culture timepoints. First, the gel is broken down by collagenase (2.5 mg/mL for 15 minutes), already delivering single cells at day 1 timepoint. For day 4 and 14, this is followed by treatment of clusters and spheroids with TrypLE (for 15 minutes) to obtain single cells. **(B)** Nuclear volumes of MCF-7 H2B-mCherry at different culture timepoints when measured in-situ and after release. For in-situ, n = 53 (day 1), 341 (day 4), 1609 (day 14); 4 gels each; N = 2. For released single cells, n = 25 (day 1), 30 (day 4), 44 (day 14); 4-6 gels each; N = 2. **(C)** Nuclear volumes of released single MCF-7 FUCCI cells from gels at different culture timepoints (grouped by cell cycle phase). n = 44 (day 1), 34 (day 4), 208 (day 14); 4-6 gels each; N = 2. **(D)** (Left) Representative slices of tomograms obtained by optical diffraction tomography (ODT) showing the refractive index (RI) of MCF-7 in gels at day 1 and 4. (Right) Histograms of the RI maps shown on the left with a Gaussian fit. **(E)** Refractive index of MCF-7 in gels at day 1 and 4. Individual dots represent average RI from whole cell or cluster. n = 51 (day 1), 28 (day 4); 4-6 gels each; N = 2-3. Data obtained in MMP-cleavable (black dots) and non-cleavable (grey dots) gels are shown. P-value obtained by likelihood ratio tests of linear mixed effects analysis in R. **(F)** Representative Brillouin frequency shift maps and corresponding brightfield images of MCF-7 in gels at day 1, 4, 14. **(G)** Box plot showing Brillouin frequency shifts of MCF-7 in gels at day 1, 4, 14. Line in the box shows the median; box shows the 25th and 75th percentile; whiskers show the 10th and 90th percentiles. n = 59 (day 1), 49 (day 4), 59 (day 14); 6 gels each; N = 3. For B, C, E: Cyan/magenta bars represent the median. For B, G: Kruskal-Wallis test with multiple comparisons (Dunn’s) was performed for statistical analysis. For E: Mann-Whitney test was performed for statistical analysis. Scale bars – (D) 25 µm; (F) 50 µm.

### Cells within spheroids increase their cytoplasmic density and lower compressibility

We suspected that the relatively rapid nuclear volume expansion upon release was due to cell volume regulation through change in intracellular water content. Therefore, we quantitatively compared the refractive index of single MCF-7 cells and small clusters (day 1 and 4) using optical diffraction tomography. By optical diffraction tomography, we obtained z-stacks of maps showing the spatial distribution of refractive indices, from which we calculated an average refractive index per object **(Fig 5D, E)**. We detected a significantly higher average refractive index for the cell clusters formed at day 4, indicating a higher mass density^47^ compared to single cells **(Fig 5E)**. We thought that these changes in cell mass density should also affect cellular compressibility. Therefore, we mapped the elastic properties of single cells and multicellular structures *in-situ* using Brillouin microscopy^48,49^ **(Fig 5F)**. We found significantly higher Brillouin frequency shifts in MCF-7 spheroids (day 14) compared to small clusters (day 4) and single cells (day 1), indicating relative increases in longitudinal moduli with cluster size increase **(Fig 5G)**, which is in agreement with previous reports^50,51^.

We excluded that the seen cell cycle changes may have accounted for overall mass density changes, since previous work has shown similar mass densities across the cell cycle^11^. Several factors might be responsible for the increased cellular mass density in spheroids compared to single cells. For instance, cells residing within dense structures might have been exposed to an increasingly crowded and hyperosmotic environment, causing an adaptive reduction of intracellular water content. Alternatively, there might be differences in cell volume regulation, e.g. due to differences in growth factor signalling and/or abundance and/or activity of ion pumps and aquaporins. Indeed, we found differential expression of various ion channels and pumps between day 1 and 4 **(Fig S10)**, like voltage gated ion (Na^+^, Ca^2+^, K^+^) channels, ligand (GABA, ATP, glutamic acid) gated ion channels and gap junctions. In summary, cells within growing cell clusters reduced their cytoplasmic and nuclear volumes, which beyond the described adjustments of their cell cycle distribution, was likely to be caused by a decrease in the cells’ relative intracellular water content, which also affected the cells’ mechanical propertie

### Invading single cells have increased nuclear volumes

We next considered the reverse scenario, where cells originating from a tumour spheroid invade the surrounding matrix – individually or as small group of invading cells. Therefore, we switched to an invasive tumour *in-vitro* model. As previously shown, Src activation can be induced in MCF10A-ER-Src cells by the addition of tamoxifen, which can lead to cell invasion in 3D culture within 72 hours^52^ **(Fig 6A)**. Quantitatively comparing invading cells and cells within the spheroid, significantly higher nuclear volumes were found for invading cells **(Fig 6B)**. Moreover, cells within invading structures of PANC1 spheroids, were found to have lower Brillouin frequency shifts compared to cells within the spheroid core **(Fig 6C, D)**. These results indicate that invading cells have increased nuclear volumes and are at the same time less stiff compared to cells within the spheroid core structure.

**Figure 6:**
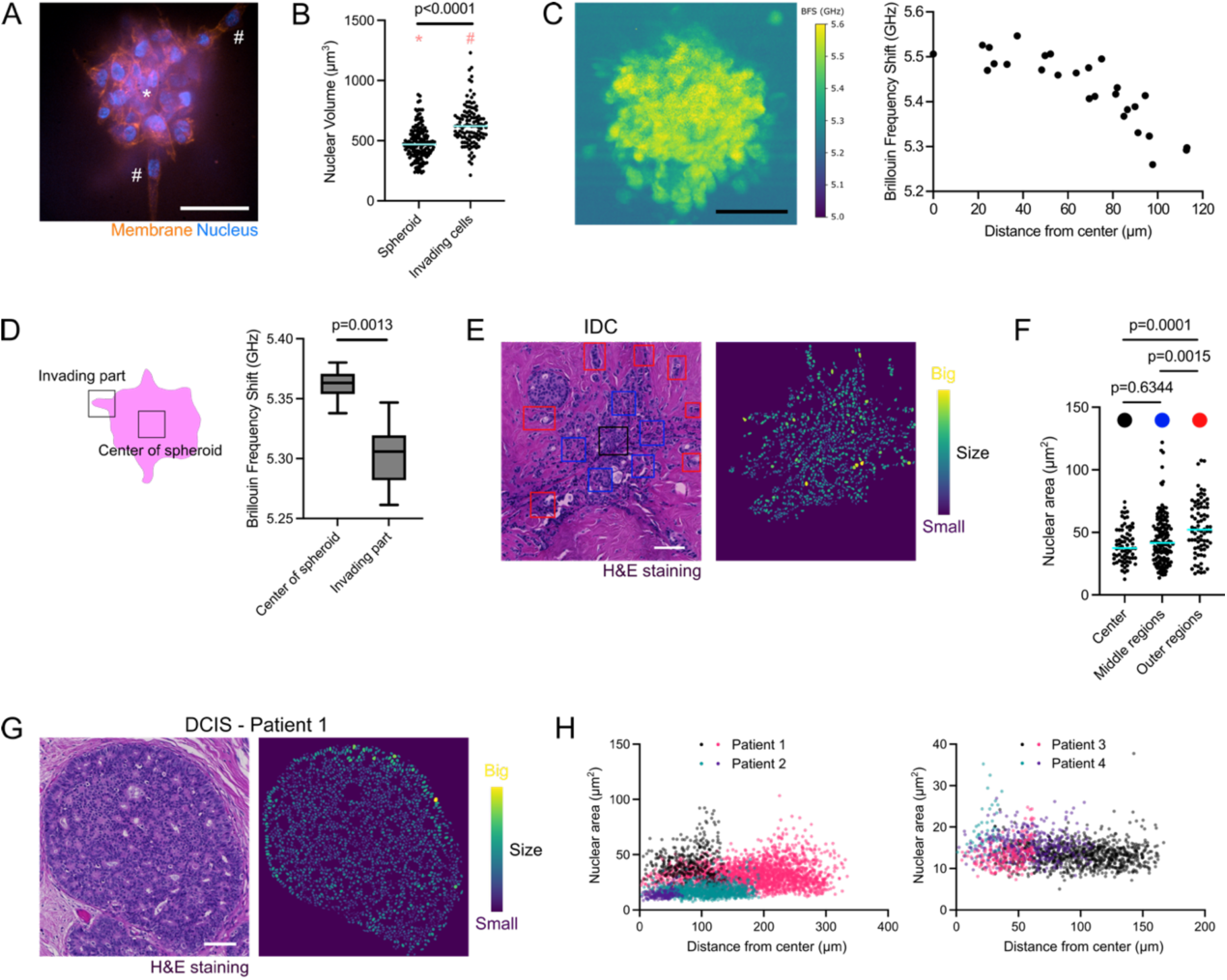
Invading cells have increased nuclear volumes compared to multicellular structures. **(A)** Representative live confocal image of MCF10A-Er-Src in compliant MMP-cleavable PEG-heparin gel stained with SiR-DNA (nucleus) and CellMask Orange (membrane) at day 8; oncogenic protein Src induced at day 5. **(B)** Scatter plot showing nuclear volumes of MCF10A-ER-Src cells within the spheroid and apparently invading out in gels at day 8. n = 171 (spheroid), 114 (invading cells); 2 gels; N = 1. **(C)** Representative Brillouin frequency shift map of PANC1 invading spheroid at day 14 in gel and corresponding Brillouin frequency shift of a defined area (∼cell size) from centre to periphery of the spheroid. n = 27; 1 spheroid. **(D)** Boxplots showing Brillouin frequency shifts from spatially resolved maps at the centre or edge of PANC1 invading spheroids in gels. Line in the box shows the median; box shows the 25th and 75th percentile; whiskers show the minimum and maximum values. n = 6 (centre of spheroid), 8 (invading part); 2 gels; N = 1. **(E)** (Left) H&E-stained section of breast IDC (invasive ductal carcinoma) with marked regions of interest. (Right) Nuclear masks from the H&E section colour coded according to size. **(F)** Scatter plot showing nuclear areas from centre, middle and outer regions of the IDC section. n = 70, 146, 78; 1 region; 1 patient. **(G)** (Left) H&E-stained sections of breast DCIS (ductal carcinoma in-situ). (Right) Nuclear masks from the H&E section colour coded according to size. **(H)** Dot plot showing nuclear areas in dependence of their distance from the centre of the DCIS tumours. n = 5314 cells over 8 tumours; 4 patients. For B, F: Cyan bars represent the median. For B, D: Mann-Whitney test was performed for statistical analysis. For F: Kruskal-Wallis test with multiple comparisons (Dunn’s) was performed for statistical analysis. Scale bars – (A, C) 50 µm; (E, G) 100 µm.

To evaluate whether these *in-vitro* findings were also present in clinical tissue samples, we then quantitated nuclear sizes in H&E-stained sections of ductal carcinoma *in situ* (DCIS) and invasive ductal carcinoma (IDC) specimens, after segmenting nuclei of epithelial cells. In contrast to our *in-vitro* model, this analysis only captures shapes and areas of nuclear slices, which differs from our 3D imaging and nuclear volume reconstruction approach on non-fixed cells in culture. The same method, however, has recently been shown to be relevant as valuable readout of cancer cell mobility^53^. In IDC, nuclei within invading structures were enlarged compared to nuclei located at the core or in between **(Fig 6E, F)**, which corroborates the observations from the *in-vitro* model where invading cells had increased nuclear volumes compared to non-invading ones. Conversely, in DCIS tumour sections **(Fig 6G)** no continuous gradient of nuclear size was observed from core to periphery **(Fig 6H)**, although at the tumour edge some nuclei displayed increased sizes. Together, these data show that volume changes observed during single to multicellular transitions can be reversed under conditions, where cells migrate out of a cell cluster, as in the case of invading tumour cells.

## Discussion

In this study, we show that there is a drastic reduction in nuclear and cell volumes as single cells form multicellular structures in 3D. We propose that this phenomenon is caused by two potentially linked mechanisms – firstly, a shift in cell cycle distribution with enrichment of smaller G1 cells over time, and in addition cell volume reduction associated with relative water content reduction, which is associated with mass density and bulk cell stiffness increase. Conversely, in an invasive tumour *in-vitro* model, where single cells invade out of a spheroid, the opposite trend is seen, where cells increase their nuclear volumes and decrease their stiffness. Importantly, this effect was seen across different cell types. Since the nuclear size reductions in invading cells were also corroborated in patient samples of breast cancer, this highlighted the relevance to study the plasticity of nuclear volume changes during tumour growth and progression.

Closely monitoring nuclear volume changes in 3D *in-vitro* cultures from day 1, to day 4 and 14, we found a progressing decrease in nuclear and indirectly also cellular volumes. The observation that over culture time more and more G1 cells accumulate, can at least partly explain the nuclear size reductions, since G1 cells have significantly lower nuclear volumes compared to G2 cells. In the same way, the higher proportion of G1 cells in free-floating spheroids might partly account for their reduced nuclear volumes. When compared to MCF-7 cells detached from 2D tissue culture plastic, representing the starting point for our 3D cultures, single cells embedded in matrix displayed a higher percentage of S/G2 cells at day 1, ∼86% versus ∼65% in 2D. This might either hint at a temporary delay in G2 or other effects causing cell cycle synchronisation between the time of matrix embedding and culture day 1. Of note, the effect of cell cycle related volume decrease from day 1 to 4 was partly reversible, since nuclear volume reductions could be rescued by treating cell clusters with the CDK1 inhibitor RO-3306, thereby enriching S/G2 cells within the clusters.

Only very few studies have directly compared cell or nuclear volumes in spheroids to single cells, and to our knowledge, none of them during growth and invasion in 3D microenvironments. For instance, Devany et al. reported up to 60% cell volume reductions for cells isolated from mature confluent MDCK monolayers compared to cells from sub-confluent 2D cultures^43^. Once the cell monolayer was mature/confluent, cells ceased to grow but continued -albeit at lower pace- to divide until inhibition of proliferation was reached. Such size-reducing divisions are, however, absent in our model, where cell growth within clusters seems to not stall along the cell cycle. Conversely, daughter cells commonly reached the size of their respective mother cell, although cell cycle progression was observed to slow down over time. There is also another reason why we rather exclude the effect of size-reducing cell divisions in our system – we find similar sized nuclei in spheroids independent of whether we started with 1, 200, 300 or 400 cells, where individual cells undergo very different numbers of divisions. We estimate that a typical day 14 spheroid in matrix contains approximately 240 cells on average. Since these cells originate from a single cell precursor, this corresponds to about 8 cell doublings. This would exceed the number of cell divisions expected to occur in the free-floating spheroids containing much less than double of the initial number of cells at day 4. Thus, the total number of divisions is unlikely to play a role in nuclear volume reductions within the forming clusters.

In contrast, two previous studies have found a minor dependence of nuclear size on the overall cell number within their spheroid cultures. In a recent study by Schmitz et al., nuclear volumes within T47D breast cancer spheroids decreased by ∼15% when comparing larger spheroids (comprising about 35000 cells) to smaller ones (about 5000 cells)^54^. These cell numbers, of course, largely exceed the numbers of cells within our spheroids. Similarly, Ong et al. reported a ∼15% decrease in nuclear volumes for colorectal tumour spheroids when cell numbers increased from ∼190 to ∼280 cells^55^, which rather matches the number of cells within our day 14 spheroids albeit in another cell type. This may suggest that the absolute cell number only has a neglectable effect on the nuclear volume within a multicellular cluster -at least beyond the point where a fairly large multicellular cluster has formed **(Fig S11)**.

In addition to the cell cycle related nuclear volume changes, we evidenced an increase in relative mass density in day 4 clusters compared to day 1 by optical diffraction tomography, even though averaging over the entire multicellular cluster may underestimate this effect, considering that these clusters also consist of ECM and solutes of lower refractive index compared to the cellular material. Since the cells and nuclei seem to rapidly swell upon cell release from multicellular structures (unlike single cells released from the matrix), we conclude that the protoplasm must have been more crowded within the clusters prior to release. Thus, two mechanisms may cause the seen nuclear/cellular volume reductions in multicellular structures, firstly an accumulation of smaller-sized G1 cells, as well as differences in cell volume regulation, potentially due to osmotic regulation.

A more crowded cytoplasm might be mechanistically linked to the given cell cycle changes. For instance, previous work has shown that hyperosmotic media can induce delays in cell cycle progression^56–58^. A possible molecular mechanism may involve higher concentrations of soluble factors that control the cell cycle, e.g. cyclins or Rb^17,43^, in shrinking cells. In line with that, longitudinal imaging of MCF-7 FUCCI cells suggested longer G1 phase lengths in the larger sized clusters at a later culture timepoint. While the addition of a CDK1 inhibitor (increasing percentage of S/G2 cells) could abolish the nuclear volume decreases that were due to a higher proportion of G1 cells, it did not fully rescue the volume decrease on day 4, suggesting that the effects on cell cycle are downstream of water-driven volume changes. Previous reports also show that changes in cell volume can affect cell cycle progression^57,59^. Unfortunately, our longitudinal imaging data was not suitable to precisely measure the nuclear volumes and cell cycle distribution in a time-resolved manner and to disentangle the temporal sequence of events. We further attempted to unravel the underlying molecular mechanisms leading to a more crowded cytoplasm, for which multiple experiments were conducted to interfere with the volume changes, e.g. sodium-hydrogen ion pumps (by EIPA), mechanosensitive ion channels (by GsMTx-4) and altering intracellular (by ionomycin and BAPTA-AM) and extracellular (by EGTA and BAPTA) calcium. Given the essential role of these ions and their pumps, we were not surprised to find increased rates of cell death when applying drugs over an extended culture time **(Fig S12)**. We therefore decided to interfere with the shorter recovery process upon single cell release at day 4 by blocking sodium-hydrogen ion pumps (by EIPA), volume regulated anion channels (by DCPIB) and aquaporin 3 (by DFP00173), but could not interfere with cell swelling upon release from the 3D multicellular structures **(Fig S12)**.

Nevertheless, we were intrigued to find in all spheroid cultures a similar minimal nuclear volume. This may suggest that there exists a lower limit of nuclear volume that is maintained by the cells’ regulatory processes. Such a lower limit on nuclear volume was also proposed by Devany et al. when studying confined 2D cultures^43^. It may be speculated that such a lower nuclear volume limit might be of physiological relevance, since crossing over such a limit could have adverse consequences on genome packaging and cause DNA damage as recently suggested^43^. With regards to a physical limit, e.g. due to crowding, it appears that cells can reduce their cellular and concomitant nuclear volumes even much further. Roffay et al. reported up to ∼90% cell volume reductions under hyperosmotic stress^34^. However, it is unlikely that cells would survive such extreme conditions for longer periods.

Unexpectedly, we did not find any substantial effect of compressive stress (up to 2.5 kPa)^40,41^ on nuclear volumes of cells within spheroids. This might be attributed to the fact that, at day 14, we might already have reached a lower nuclear volume limit as discussed above. Previous work from our lab has reported an increased percentage of G1 cells (by ∼10%) in MCF-7 spheroids under compressive stress (for spheroids in stiff compared to compliant MMP-cleavable hydrogels)^40^, which would be expected to cause a minor decrease in mean cellular and nuclear volumes. While we could not detect significant decreases in median nuclear volumes in stiffer matrices here, it is possible that the associated compressive stress might have affected the respective cell volumes, which we could not verify *in-situ* though. For instance, a previous study by Vahala et al. showed that when tumour spheroids are cultured in stiffer gels, their cell volumes decreased while nuclear volumes remained rather similar^60^. Here, we have only quantitatively assessed nuclear volumes, while whole cell volumes were only indirectly measured for multicellular structures (cell volumes could only be reliably measured for *in-situ* or released single cells) **(Fig 2G)**. Although it would be ideal to directly measure both nuclear and cytoplasmic volumes in our model, 3D segmentation of cell volumes within dense multicellular structures remains a challenge.

We further observed that cells invading out of the spheroids had larger nuclear volumes. This finding concords with previous reports by Han et al. showing cell swelling at the invasive front of mammary epithelial (MCF10A) spheroids^37^. Han et al. have attributed the larger sized nuclei of invading cells to a continuous volume gradient formed by intercellular water flows (due to compressive stress) from the spheroid core towards the periphery^37^. Such a gradient, however, cannot be confirmed by our model. Even though our experiments bear similarities (i.e. MCF-10A clusters within hydrogels), we could not find any consistent radial gradients in nuclear volumes or cell-cell distances for MCF-7 and MCF10A spheroids **(Fig S13)**. Our findings are also in good agreement with other reports that did not observe any significant variations in nuclear volumes or cell-cell distances across the spheroid radius ^26,61,54^. For instance, Ong et al. recently detected only 5% difference in nuclear volumes of cells at the core and periphery of similar sized HCT116 (colon cancer) spheroids^55^. Similarly, Monnier et al. found gradients in cell volumes only when exposing CT26 (colon cancer) spheroids to compressive stress induced by osmotic pressure increase, but not under isotonic conditions^61^, which reflects previous reports by Delarue et al. on HT29 (colon cancer), CT26 (mouse colon cancer) and BC52 (breast cancer) spheroids^26^. Thus, instead of intercellular water flow and the establishment of a cell volume gradient across the multicellular structure, we rather hypothesise that cell volume regulation is dependent on the multicellular context of the cells and changes when cells disconnect from their neighbours. Larger sized cells at the edge of the spheroid that have lost connection to some of their neighbouring cells, might represent a reversion of the phenomenon seen when single cells form multicellular structures. Despite possible adaptions in cell cycle, cells might swell, once they lose the connection to their neighbouring cells and migrate out of the cluster, as observed upon releasing single cells from multicellular structures. Thus, swelling might rather be a consequence than cause of cell dissemination from the clusters.

These invading cells with increased nuclear volumes were in our case also characterised by a lower Brillouin frequency shift compared to cells residing within the cluster, suggesting a more compliant cellular phenotype, which is in line with Han et al. reports showing a lower apparent modulus of invading cells compared to cells in the spheroid core^37^. These findings highlight how tightly cell volume and mechanical properties are coupled^30^ and relate to the function of cancer cells. A softer cellular phenotype is typically associated with increased invasive potential of cancer cells, as shown for multiple cancer cell lines but also primary cells^62,63^. While such mechanical chances can be related to cytoskeletal changes due to oncogene activation or EMT programs, they can also be part of an adaptive cellular response, e.g. due to confinement as previously suggested^64^, but possibly also to the multicellular context as shown here. The relationship of the cell mechanical phenotype and its invasive potential is probably much more complex though. Softening of cells and their nuclei due to swelling might not be advantageous for cells navigating through tight tissue spaces, which may rather benefit from smaller and more deformable nuclei^65^. Beyond that, cancer cells are thought to increase their survival chances under shear within the blood circulation by forming clusters on their metastatic route^66^. Although speculative, the adoption of a more compact structure within such clusters and concomitant cellular stiffening may further contribute to this effect. Our findings also have some practical implications for researchers working with 3D models. To form 3D models, cells are commonly aggregated in suspension or embedded into a 3D matrix to build more physiologically relevant models, for instance to study drug response. The volume and cell cycle changes we reported here might be very relevant in such cases, since they are likely to influence multiple cellular processes including cell signalling and gene expression^67,68^.

In summary, we show that cells dynamically adjust their volumes and cell cycle during single to multicellular state transitions, which in the context of a tumour cell, could have important implications during tumour growth and invasion. Still, there are several mechanisms that remain to be explored in more detail, for which the presented 3D model can be used. Importantly, the shown volume plasticity appears not to be a tumour cell specific phenomenon, but may also be of relevance for other physiological and pathological processes where cells switch between single and multicellular cell states.

## Methods

### Cell culture

MCF-7 cells were obtained from the “Deutsche Stammsammlung für Mikroorganismen und Zellkulturen” (DSMZ). Cells were cultured in advanced RPMI 1640 supplemented with 10% heat-inactivated (30 minutes at 56°C) fetal bovine serum (FBS), 1% MEM non-essential amino acids, 10 µg/mL human insulin and penicillin (100 U/mL)-streptomycin (100 µg/mL). PANC1 cells were kindly provided by Prof. Dr. med. Christoph Kahlert, UKD, Dresden. Cells were cultured in DMEM-GlutaMAX supplemented with 10% FBS and penicillin (100 U/mL)- streptomycin (100 µg/mL). MCF10A H2B-GFP cells were kindly provided by Dr. Helen Matthews, University of Sheffield. Cells were cultured in DMEM/F12-GlutaMAX supplemented with 5% horse serum, 20 ng/mL EGF, 0.5 µg/mL hydrocortisone, 100 ng/mL cholera toxin, 10 µg/mL human insulin and penicillin (100 U/mL)-streptomycin (100 µg/mL). MCF-7 FUCCI cells were kindly provided by Costanza Giampietro, EMPA, ETH Zurich. Cells were cultured in DMEM-GlutaMAX supplemented with 10% heat-inactivated FBS and penicillin (100 U/mL)-streptomycin (100 µg/mL). HeLa FUCCI cells were kindly provided by A. Ferrari, ETH Zurich. Cells were cultured in DMEM supplemented with 10% FBS and penicillin (100 U/mL)- streptomycin (100 µg/mL). MCF10A-ER-Src cells were kindly provided by Prof. Kevin Struhl, Harvard Medical School. Cells were cultured in the same media as MCF10A H2B-GFP except for two differences – phenol free media and horse serum was charcoal stripped (protocol for which can be found here https://genome.ucsc.edu/encode/protocols/cell/human/MCF-10A_Struhl_protocol.pdf). Cells were cultured in 2 µg/mL puromycin for a couple of passages, followed by 0.5 µg/mL puromycin to remove any cells that might lose the ER-Src plasmid.

Before starting 3D cultures, cells were grown in standard T25 culture flasks and sub-passaged 2–3 times per week. For cell detachment, TrypLE was used. All cell culture reagents were from Thermofisher unless otherwise stated.

### Generation of MCF-7 H2B-mCherry and PANC1 H2B-mCherry cells

2 × 10^5^ MCF-7/PANC1 cells were seeded per well in a 24-well plate. H2B-mCherry plasmid (#20972) transfection was performed using Lipofectamine 2000 based on manufacturer’s protocol. After 48 hours, the well with best transfection efficiency (as seen by fluorescence) was transferred to a 6-well plate. 800 µg/mL (for MCF-7) and 1200 µg/mL (for PANC1) G418 was added to the cells next day for selection of cells that had taken up the plasmid. For about 2 weeks, fresh media was exchanged with G418. Then, single cell clones were made in a 96-well plate from these cells. 5-6 single cell colonies positive for fluorescence were pooled together, expanded and frozen. The transfected cell lines were cultured in the same media as the parent cell line.

### Bioengineered hydrogel-based 3D cell cultures

PEG and heparin-maleimide precursors were synthesized as previously described^39^. Cells were detached from a T-25 flask with TrypLE, resuspended in cell culture medium, centrifuged and resuspended in PBS. Then, cells were mixed with freshly prepared heparin-maleimide (MW∼15000) solution and RGD peptide (final 3.57 µg/µL), at a density of 800 cells/µL. The final heparin-maleimide concentration was 1.5 mM. PEG precursors with either MMP-cleavable peptide (MW∼15000-15500) or not (MW∼10000), were reconstituted with PBS at a concentration of 2.25 mM (for stiff gels) along with 1 M HCl (∼1:270 – 1-part HCl to 270-parts PEG solution) for a pH of ∼6. Then it was placed into an ultrasonic bath (Merck) for 1 minute (medium intensity). This PEG solution was used to prepare stiff gels (15-25 kPa Young’s modulus). Compliant gels (1-3 kPa Young’s modulus) were prepared by diluting the stock solution of PEG precursors (∼1.05 mM). Based on the functionalisation of PEG, we were able to make compliant and stiff MMP-cleavable and non-cleavable gels. To prepare PEG-heparin hydrogel droplets, heparin-cell suspension was mixed in chilled microcentrifuge tubes with the same amount of PEG solution using a low binding pipette tip. Then, a 20 µL drop of the PEG-heparin-cell mix was pipetted onto Sigmacote-treated (hydrophobic) glass slides. Gel polymerization started immediately. Hydrogels were gently detached from the glass surface after 7 minutes using a razor blade and transferred into a 24-well plate supplemented with 1 mL cell culture medium. Cell culture medium was exchanged every 3 days. Unless mentioned, experiments were performed with cells cultured in compliant MMP-cleavable gels.

### Cell release from hydrogel cultures

At each culture timepoint, hydrogels were treated with 2.5 mg/mL collagenase for 15 minutes at 37 °C. At 10 minutes, the solution was additionally pipetted to mechanically break the gels as well. Released cells or multicellular structures were centrifuged at 180 × g for 2 minutes. The collagenase was removed and cells were washed with PBS. At day 1, single cells were then resuspended in CO_2_ independent media and were ready for imaging. At day 4 and 14, multicellular structures were treated with TrypLE for 15 minutes at 37 °C (with a similar intermediate pipetting step at 10 minutes). The broken cells were also washed with PBS once, before resuspending them in CO_2_ independent media, to be used for imaging.

### Quantification of hydrogel mechanical properties by AFM (atomic force microscopy)

After hydrogel preparation, they were immersed in PBS and stored at 4 °C until mechanical characterization on the following day. After equilibrating them at room temperature for 1 hour, gels were put on 35 mm plastic dishes and held down by a metal ring with strings (ALA Scientific Instruments, NY, USA). A Nanowizard IV (JPK Instruments, Berlin, Germany) was used to probe the gel stiffness. Arrow T1 cantilever that had been modified with a polystyrene bead of 5 or 10 μm diameter using epoxy glue, was calibrated using the thermal noise method implemented in the AFM software (Version 6.1.159, NanoWizard Control Software, JPK Instruments/Bruker, Berlin, Germany). Hydrogels were probed at room temperature in PBS using a speed of 5 μm/s and a force setpoint of 2-6 nN (based on hydrogel stiffness) in order to obtain indentation depths of 0.5-1 µm. Force distance curves were processed using the JPK data processing software (Version 6.1.159, NanoWizard Control Software, JPK Instruments/Bruker, Berlin, Germany) using the Hertz/Sneddon model for a spherical indenter, assuming a Poisson ratio of 0.5^69,70^.

### Staining of frozen sections

MCF-7 cells were cultured in hydrogels with fluorescently labelled heparin and fixed with 4% paraformaldehyde for 45 minutes on day 14. Then, hydrogels were transferred into 30% of sucrose/PBS for 1 hour, and thereafter embedded into optimum cutting temperature cryo-embedding compound (O.C.T., Thermofisher) and snap frozen in liquid nitrogen. 3D samples were sectioned in 8–10 µm thick cryosections with the Microm HM560 Cryostat (Thermofisher). Antigen retrieval was performed by immersing slides into citrate buffer (0.1 M, pH 6.0)/ 0.05% Triton X-100 (Sigma) for 10 minutes. Then slides were incubated for 10 minutes in 0.1% Triton X-100/PBS and blocked for 1 hour in 2.5% BSA/PBS. Slides were incubated with CK8/18 (Dianova) primary antibody in 2.5% BSA/PBS overnight at 4–6 °C in a humidified chamber. After washing the glass slides with PBS, slides were incubated for 2 hours with secondary antibody (Cy5-conjugated anti-mouse/rabbit IgG, Dianova), Phalloidin-TRITC (Sigma) and DAPI (Sigma). Finally, the slides were washed with PBS and mounted after a short wash in ddH_2_O with mounting medium (Thermofisher). Samples were imaged with a LSM700 confocal microscope (Zeiss) using a 40× objective (C-Apochromat, Zeiss).

### Imaging of fixed single cells and multicellular structures

On day 1, 4 and 14, cultures were fixed with 4% paraformaldehyde in PBS for 45 minutes, followed by permeabilization in 0.2% Triton X-100 for 30 minutes. Spheroids were stained for 4 hours with 5 µg/mL DAPI and 0.2 µg/mL Phalloidin-TRITC in 1% BSA in PBS with 0.1% Triton X-100. Hydrogels were stored in PBS in the fridge. Single cells and multicellular structures were imaged with a LSM780 confocal microscope using a 40× objective (Zeiss C-Apochromat, White Plains, NY, USA).

### Brillouin microscopy

Brillouin maps were obtained on a setup previously described^41,71^. The setup was a custom-built confocal Brillouin microscope employing a two-stage VIPA (virtually imaged phased array) spectrometer. The setup featured frequency-modulated diode laser with a wavelength of 780 nm whose frequency was stabilized to the D2 transition of Rubidium 85. To suppress background light created by amplified spontaneous emission, a Bragg grating and a Fabry– Pérot interferometer in a two-pass configuration was used. Imaging was done with a 20×/0.5 air or 40×/0.95 air objective and the sample temperature was controlled at 37°C by a petri dish heater. The gels were mounted on a 35 mm glass bottom disk with CO_2_ independent media and held down by custom made rings with a series of strings. Brillouin maps were analysed either by a MATLAB software ‘BrillouinEvaluation’ or a python software ‘BMicro’ (https://github.com/BrillouinMicroscopy). Brillouin maps were segmented in ‘Impose’ (https://github.com/GuckLab/impose) or Fiji (using maps exported as tiff by BMicro). Median was calculated from all BFS values within the region of interest (corresponding to a single cell or multicellular structure).

### Live spinning disk confocal microscopy

Cell membrane of single cells and multicellular structures was stained with CellMask Deep Red or Orange (concentration of 1:1000 in culture media) by incubation for 1 hour at 37 °C and washing three times with PBS. For MCF10A-ER-Src cells which did not have the nucleus fluorescently labelled, SiR-DNA was used for staining the nucleus. Cultures were incubated with 1:1000 SiR-DNA for 6 hours at 37 °C. The ibidi 4 or 8-well glass bottom µ-slide was used. Imaging was done with a 40×/0.95 air objective on a Nikon Ti-E confocal microscope with spinning disk and a heated stage (37 °C) with CO_2_ independent media. Resolved z-stacks were obtained with 0.5 µm spacing in z.

### Cell and nuclear segmentation

The following steps were done to obtain the cell and nuclear volume once the z-stacks were obtained on the fluorescence spinning disk confocal microscope:

- Background subtraction – using rolling ball method (radius=50pixels) in Fiji (1.54f)^72^.
- Cropping – images were cropped in Fiji to remove empty space around the cells or multicellular structures which would otherwise slow down processing.
- Ground truth data generation – nuclear segmentation required training of a StarDist^73–75^ model for which ground truth data was needed. First, Cellpose GUI (v2.2)^76,77^ was used to modify their pretrained ‘nuclei’ model using the human-in-loop training on a few 2D slices of the nuclear channel. This new model was then used to segment the nuclei in two spheroids using the Cellpose plugin in Napari (0.4.17)^78^. Using the labels layer in Napari, the wrong labels were corrected and missing labels were manually added.
- Training model in StarDist – the ground truth data was used to train a StarDist model with default parameters.
- Performing nuclear segmentation – with the custom StarDist model, segmentation was either done using the StarDist plugin in Napari or with the ZeroCostDL4Mic StarDist3D notebook on GoogleColab^79^. Incorrect masks were removed in Napari.
- Whole object segmentation for cell volume – Limeseg^80^, a plugin in Fiji, was used on the membrane channel to either segment single cell volume (using ‘sphere seg’) or segment volume of whole multicellular structure (using ‘skeleton seg’), which was then divided by the number of nuclei to get the average cell volume.

### Analysis of segmented objects

Once the nuclei were segmented through StarDist and their 3D masks were obtained, their volumes were quantified by the 3D Volume feature under the 3DSuite Fiji plugin. As for the cells or whole objects segmented with Limeseg, the plugin in Fiji measured and reported the volume. The sphericity of the 3D nuclei masks was measured by the 3D Compactness feature under the 3DSuite Fiji plugin. For the spatial analysis in spheroids, the centre of the nuclei was determined by 3D Centroid in 3DSuite. After obtaining the centroid of the spheroid by segmenting the whole object with Limeseg, the distance of each segmented nuclei from the centre was calculated. The distance between each nucleus and its closest neighbour was also measured with 3DSuite, with the 3D Distances feature.

### Numerical simulations for volume and cell cycle variation with multicellularity

We performed mechanistic simulations of growing and dividing spherical cells in two dimensions, starting from a single cell. All results shown were averaged over 1000 simulation runs. Exponential growth of cells was prescribed with a growth rate randomly chosen for each cell at its birth, following the log-normal distribution ℒ𝒩(−3.8139, 0.2738). The used parameters were fitted from measured cell cycle times, implying an average rate of *e*^-3.8139^ = 0.0221/h.

The G1 state was defined by the cell size (area in 2D) being smaller than 1.26 times the initial size. This value reflected the experimentally observed cycle times (≈10h in G1, ≈20h in G2, at days 2 to 4) under the exponential growth. Once a cell doubled its size, it divided into two daughter cells of equal size. The division plane was randomly chosen. To avoid overlap of cells, a repulsive force was implemented such that cells pushed their neighbours away. As the amount (*n*) of cells in the cluster grew, the growth rate gradually decreased in G1 state by a factor of 1/(1 + 0.015*n*) to mimic the extended G1 cycle times measured for larger populations (≈23h in G1, ≈20h in G2, at days 8 to 10).

Treatment with RO-3306 was included in the model by stopping cell division after day 3, whereupon cells were halted in G2 and growth of the affected cells was stalled. Simulations are implemented and run in Matlab R2022a.

### Optical diffraction tomography

The refractive index (RI) distribution of single cells and multicellular structures in hydrogels was determined using optical diffraction tomography (ODT). This method employed Mach-Zehnder interferometry to acquire multiple complex optical fields from various incident angles, following previously established protocols^11,49^. A frequency-doubled Nd-YAG laser (λ = 532 nm, Torus 532, Laser Quantum Ltd, UK) was coupled into a single-mode fiber and split into two beams by a 2×2 fiber coupler. One beam served as the reference and the other beam directed onto the sample stage of a custom-made inverted microscope through a tube lens (f = 175 mm) and a water-dipping objective lens (NA = 1.0, 40×, Carl Zeiss AG). The diffracted light from the sample was collected using a high numerical aperture objective lens (NA = 1.2, 63×, water immersion, Carl Zeiss AG). The diffracted beam interfered with the reference beam at the image plane, producing a spatially modulated hologram, which was recorded using a CCD camera (FL3-U3-13Y3M-C, FLIR Systems Inc.). To construct a 3D RI tomogram, the sample was illuminated from 150 different angles, which were scanned by a dual-axis galvanomirror (GVS012/M, Thorlabs Inc.) located at the samples conjugate plane. From the measured holograms, complex optical fields scattered by the sample were retrieved using a Fourier transform-based field retrieval algorithm^81^. The 3D RI distribution of the sample was reconstructed from these retrieved fields via the Fourier diffraction theorem, employing the first-order Rytov approximation^82,83^. Additional details on the tomogram reconstruction process are available in previous studies^84^. The MATLAB script for ODT reconstruction can be accessed at: https://github.com/OpticalDiffractionTomography/ODT_Reconstruction. To be able to perform ODT on single cells and multicellular structures, gels were sandwiched between 35 mm glass bottom dish and Sigmacote-treated 13 mm circular coverslips. Once the gels polymerized, the dish was filled with media and the circular coverslip was gently pushed off and removed. This gave rise to a flat gel on the dish which was compatible with the low working distance of the ODT setup. The analysis scripts resulted in tiff files of the tomograms. The single cells and small clusters were segmented in Cellpose (2.2) in 3D and resulting masks were used to obtain an average RI value using the ‘3D Intensity Measure’ function in 3DSuite in Fiji.

### RNA isolation, sequencing and analysis

RNA isolation – released single cells (day 1) and multicellular structures (day 4) were obtained by treatment of compliant and stiff MMP-cleavable hydrogels with 2.5 mg/mL collagenase for 15 minutes at 37 °C. RNA was isolated using Trizol using the manufacturer’s protocol. RNA concentration and quality were assessed by Bioanalyzer (Agilent 2100). For library preparation, mRNA was isolated from 90 ng total RNA by poly-dT enrichment using the NEBNext Poly(A) mRNA Magnetic Isolation Module (NEB) according to the manufacturer’s instructions. Samples were then directly subjected to the workflow for strand-specific RNA-Seq library preparation (Ultra II Directional RNA Library Prep, NEB). For ligation, NEB Next Adapter for Illumina (0.15 µM) of the NEB Next Multiplex Oligos for Illumina Kit were used. After ligation, adapters were depleted by an XP bead purification (Beckman Coulter) adding the beads solution in a ratio of 0.9:1 to the samples. Unique dual indexing was done during the following PCR enrichment (14 cycles) using amplification primers carrying the same sequence for i7 and i5 index (i5: AAT GAT ACG GCG ACC ACC GAG ATC TAC AC NNNNNNNN ACA TCT TTC CCT ACA CGA CGC TCT TCC GAT CT, i7: CAA GCA GAA GAC GGC ATA CGA GAT NNNNNNNN GTG ACT GGA GTT CAG ACG TGT GCT CTT CCG ATC T). After two more XP bead purifications (0.9:1), libraries were quantified using the Fragment Analyzer (Agilent) using the NGS Kit. Libraries were sequenced on an Illumina NovaSeq 6000 in 100 bp paired-end mode to a depth of 40-60 million fragments per library. For obtaining gene counts, FastQC (http://www.bioinformatics.babraham.ac.uk/) was used to perform a basic quality control of the resulting sequencing data. Fragments were aligned to the human reference genome hg38 with support of the Ensembl 98 splice sites using the aligner STAR (2.7.10b)^85^. Counts per gene and sample were obtained based on the overlap of the uniquely mapped fragments with the same Ensembl annotation using featureCounts (v2.0.1)^86^. Gene set and differential expression analysis – To get gene sets, GSEA (gene set enrichment analysis) (https://www.gsea-msigdb.org/gsea/index.jsp) was used for all the genes against the MSigDB database (H, C2, C3, C4, C5, C6, C8 collections). Gene sets relevant to a certain process were searched with the relevant keyword (e.g. ‘cell_cycle’ was the keyword used to get all gene sets involved in cell cycle). Gene sets with NOM p-val < 0.05 and FDR q-val < 0.25 were plotted or further used for leading edge analysis. To plot specific genes for the enriched gene sets, leading edge analysis was performed in GSEA to obtain a list of genes which were present in majority of gene sets. z-scores for these genes were calculated from the normalized counts obtained through DESeq2 in RStudio (2024.04.0+735) from the raw gene counts (http://www.bioconductor.org/packages/release/bioc/html/DESeq2.html). DESeq2 was also used for differential gene expression analysis and allowed for the calculation of fold change in the expression of genes between the conditions. An online database for genes for cellular pumps/channels was used (https://www.guidetopharmacology.org/GRAC/IonChannelListForward?class=VGIC). Differentially expressed genes with padj < 0.05 and log2FoldChange > 0.5 or < −0.5 which were present in the database were plotted.

### Coarse longitudinal imaging with confocal microscopy

Coarse longitudinal imaging was done in two scenarios – in one, single cells were imaged every 12 hours from day 0 to day 4 and another where the same cells were imaged at day 1 and then at day 4. For both these cases, gels were formed on a 35 mm glass bottom dish with a grid etched on them (ibidi Grid-500). Initially, the xy position of each cell on the grid and the z position from the microscope was noted. These xyz positions allowed for the same entity to be tracked over multiple imaging rounds. Imaging was done with a 40×/0.95 air objective on a Nikon Ti-E confocal microscope with spinning disk and a heated stage (37 °C) with CO_2_ independent media. Resolved z-stacks were obtained with 0.5 µm spacing in z. Between imaging sessions, the samples were normally incubated in the 37 °C, 5% CO2 incubator with culture media.

### Resolved timelapse imaging with light sheet microscopy

Light sheet imaging was performed using Swept Confocally-Aligned Planar Excitation (SCAPE) microscopy (ASI, custom single-objective SCAPE, version CB1v0), equipped with a Leica HC PL APO 40×/1.10 NA objective and twoTeledyne Kinetix22 cameras. Samples were maintained at 37 °C and 5% CO₂ in a heated chamber (Okolab, H301-NIKON-NZ100/200/500-N) fitted with a 35 mm #1 glass-bottom dish holder (Okolab, 1×35-M). Environmental conditions were regulated using the UNO-T-H-CO2 all-in-one controller with CO₂/air mixer (Okolab), and the objective was heated using the OBJ-COLLAR-2532 (Okolab).

Angular distortion resulting from the angled light sheet was corrected using a shearing operation based on an affine transformation, implemented via the py-clesperanto library (https://github.com/clEsperanto/pyclesperanto). Time-lapse imaging was conducted with a temporal resolution of 30 minutes and a voxel size of 0.178 × 0.178 × 0.5 µm.

For measuring the cell cycle phase lengths, fluorescence intensity profiles were plotted over time for Cdt1 and Geminin, either from 3D segmented nuclei (using StarDist as described earlier) or by manually selecting ROIs over nuclei in a maximum intensity 2D projection. G1 length was calculated from birth till the peak of Cdt1 intensity was reached. S/G2 length was calculated from the beginning of steady increase in Geminin intensity till nuclear envelope breakdown. If the cell was captured from birth or till nuclear envelope breakdown, this resulted in the absolute quantification of G1 and S/G2 length, respectively. If not, the length was reported as being greater than a certain duration. For measuring the growth rate, nuclear segmentation of nuclei (from StarDist) between day 1 to 4 was taken. The nuclear volume over time, irrespective of cell cycle phase, was fitted to an exponential growth equation in Prism.

### Analysis of patient-derived DCIS and IDC tissue slices

FFPE (formalin fixed paraffin embedded) samples were prepared by the Tumour and Normal Tissue Bank (TNTB) of the BioBank Dresden and originate from the archive of the Institute of Pathology (EK59032007) (Prof. Dr. rer. nat. Jan Kuhlman) of the University Hospital Dresden. H&E staining was performed by the Institute of Pathology of the University Hospital Dresden. Whole slide images were acquired on an Axioscan Z.1 (Carl Zeiss) microscope of the CMCB light microscopy facility, a Core Facility of the CMCB Technology Platform at TU Dresden, using a Plan-Apochromat 10×/ 0.45 (Carl Zeiss) objective and a HV-F202SCL (Hitachi) colour camera. The nuclear area from H&E images of DCIS and IDC tumours were obtained though the StarDist plugin in Fiji. The versatile H&E model was used with default parameters. The resulting masks were cleaned up in Napari.

### Forming free-floating aggregates

To form free-floating aggregates, cells were detached from a T-25 flask with TrypLE, resuspended in cell culture medium, centrifuged down and resuspended again in cell culture media. 200, 300 and 400 cells were added per well to a 96-well suspension U-bottom plate in 100 µL (day 0). The plate was centrifuged at 290 × g for 3 minutes. The next day (day 1), 100 µL media containing 6 µg/mL rat tail collagen I was added to each well (media kept at room temperature). The plate was centrifuged at 100 × g for 3 minutes. On day 4, the self-aggregating spheroids were ready to be imaged. Spinning disk confocal microscopy was performed as described above.

### MCF10A-ER-Src activation for invading spheroids

Single MCF10A-ER-Src cells were embedded in compliant MMP-cleavable gels as described above. The cells were allowed to form spheroids till day 5. At day 6, 1 µM tamoxifen was added to the cultures to transform the cells. On induction of the Src oncogene, the spheroids started to form invading structures and were imaged with SiR-DNA and CellMask Orange (as described above) on day 8.

### Statistical analysis

GraphPad Prism (10.4.2) was used to plot data and to perform statistical tests. A two-tailed significance level of 5% was considered statistically significant (p < 0.05). As indicated in the figure legends, for pairwise comparisons, a Mann–Whitney test was used since most datasets were not normally distributed. In the case of more than two groups, a Kruskal–Wallis with multiple comparisons (Dunn’s) was chosen. Linear regression was performed to look at correlation between two variables. Linear mixed effects analysis on ODT data was performed using RStudio (Version 2023.06.2+561) and lme4 (Bates, Maechler & Bolker, 2012). As fixed effects, we considered culture day (1 or 4), as random effect we had intercepts for gel type (MMP-cleavable or non-cleavable). P-values were obtained by likelihood ratio tests of the full model with the effect in question against the model without the effect in question.

## Acknowledgements

We thank the Histology Facility, CMCB, TU Dresden for help with the cryo-sectioning. We thank the light microscopy facility, a Core Facility of the CMCB Technology Platform at TU Dresden, and the Biophysical Core facility (MPI-CBG), especially Dr. Ellen Geibelt, Dr. Hella Hartmann, Dr. Nazar Oleksiievets and Dr. Jens Ehrig, for excellent technical support. We also thank Pierre DeMarinis for support with imaging and analysis. Bulk RNA-sequencing and initial analysis for reads was performed by the Dresden-concept Genome Centre, by Grit Mehnert and Mathias Lesche. Sevina Dietz and DcGC helped with the bioanalyzer to measure RNA concentration and integrity. We also thank Prof. Ingmar Glauche for statistical advice, and Bruker JPK, Berlin and CellSense, Berlin for technical support regarding AFM and Brillouin microscopy, respectively. We thank Prof. Jan Kuhlman for support with tissue sections. DCIS and IDC patient sections were provided and stained by the BioBank Dresden, a core facility of the Medical Faculty Dresden/Technical University Dresden and a National Centre of Tumour Diseases Dresden resource. Vaibhav Mahajan and Anna Taubenberger are part of the Mildred Scheel Nachwuchszentrum Dresden IP3 funded by the German Cancer Aid. The project was further supported by the German Research Society (DFG) through TA 751/4-1 and AL 1705/11-1.

## Author contributions

Conceptualization, A.V.T., V.M.; methodology, V.M., B.L., K.K., M.M., S.A., R.S., A.V.T.; software, V.M., B.L, K.K., M.M., S.A., R.S.; validation, V.M.; formal analysis, V.M., K.G.B., A.G., V.A.K, S.A., T.A.; investigation, V.M., K.G.B., A.G., T.B., M.M., S.A., A.V.T.; resources, C.W., A.H., A.V.T.; data curation, A.V.T., V.M.; writing—original draft preparation, V.M., A.V.T.; writing—review and editing, all authors; visualization, A.V.T., V.M.; supervision, A.H., A.V.T., R.S.; funding acquisition, C.W., A.H., S.A., A.V.T. All authors have read and agreed to the published version of the manuscript.

## References

1. Tscherner, A. K., Macaulay, A. D., Ortman, C. S. & Baltz, J. M. Initiation of cell volume regulation and unique cell volume regulatory mechanisms in mammalian oocytes and embryos. J. Cell. Physiol. 236, 7117–7133 (2021).

2. Zucker, R. M., Whittington, K. & Price, B. J. Differentiation of HL-60 cells: Cell volume and cell cycle changes. Cytometry 3, 414–418 (1983).

3. Urbanska, M. et al. Single-cell mechanical phenotype is an intrinsic marker of reprogramming and differentiation along the mouse neural lineage. Development 144, 4313–4321 (2017).

4. Lengefeld, J. et al. Cell size is a determinant of stem cell potential during aging. Sci. Adv. 7, eabk0271 (2021).

5. Laforest, S., Labrecque, J., Michaud, A., Cianflone, K. & Tchernof, A. Adipocyte size as a determinant of metabolic disease and adipose tissue dysfunction. Crit. Rev. Clin. Lab. Sci. 52, 301–313 (2015).

6. Davies, D. M., van den Handel, K., Bharadwaj, S. & Lengefeld, J. Cellular enlargement -A new hallmark of aging? Front. Cell Dev. Biol. 10, (2022).

7. Manohar, S. & Neurohr, G. E. Too big not to fail: emerging evidence for size-induced senescence. FEBS J. 291, 2291–2305 (2024).

8. Ginzberg, M. B., Kafri, R. & Kirschner, M. On being the right (cell) size. Science (2015) doi:10.1126/science.1245075.

9. Schmoller, K. M. The phenomenology of cell size control. Curr. Opin. Cell Biol. 49, 53–58 (2017).

10. Cadart, C., Venkova, L., Piel, M. & Cosentino Lagomarsino, M. Volume growth in animal cells is cell cycle dependent and shows additive fluctuations. eLife 11, e70816 (2022).

11. Kim, K. & Guck, J. The Relative Densities of Cytoplasm and Nuclear Compartments Are Robust against Strong Perturbation. Biophys. J. 119, 1946–1957 (2020).

12. Cadart, C. et al. Size control in mammalian cells involves modulation of both growth rate and cell cycle duration. Nat. Commun. 9, (2018).

13. Chandler-Brown, D., Schmoller, K. M., Winetraub, Y. & Skotheim, J. M. The Adder Phenomenon Emerges from Independent Control of Pre-and Post-*Start* Phases of the Budding Yeast Cell Cycle. Curr. Biol. 27, 2774–2783.e3 (2017).

14. Rhind, N. Cell-size control. Curr. Biol. 31, R1414–R1420 (2021).

15. Ginzberg, M. B. et al. Cell size sensing in animal cells coordinates anabolic growth rates and cell cycle progression to maintain cell size uniformity. eLife 7, e26957 (2018).

16. Liu, X., Yan, J. & Kirschner, M. W. Cell size homeostasis is tightly controlled throughout the cell cycle. PLOS Biol. 22, e3002453 (2024).

17. Zatulovskiy, E., Zhang, S., Berenson, D. F., Topacio, B. R. & Skotheim, J. M. Cell growth dilutes the cell cycle inhibitor Rb to trigger cell division. Science 369, 466–471 (2020).

18. Ma, Y. & Edgar, B. A. CDK4: Linking cell size to cell cycle control. Dev. Cell 56, 1695–1696 (2021).

19. Chadha, Y., Khurana, A. & Schmoller, K. M. Eukaryotic cell size regulation and its implications for cellular function and dysfunction. Physiol. Rev. 104, 1679–1717 (2024).

20. Kellogg, D. R. & Levin, P. A. Nutrient availability as an arbiter of cell size. Trends Cell Biol. 32, 908–919 (2022).

21. Saxton, R. A. & Sabatini, D. M. mTOR Signaling in Growth, Metabolism, and Disease. Cell 168, 960–976 (2017).

22. Hoffmann, E. K., Lambert, I. H. & Pedersen, S. F. Physiology of cell volume regulation in vertebrates. Physiol. Rev. 89, 193–277 (2009).

23. Yancey, P. H. Organic osmolytes as compatible, metabolic and counteracting cytoprotectants in high osmolarity and other stresses. J. Exp. Biol. 208, 2819–2830 (2005).

24. Clark, A. G. & Paluch, E. Mechanics and regulation of cell shape during the cell cycle. Results Probl. Cell Differ. 53, 31–73 (2011).

25. Fischer-Friedrich, E., Hyman, A. A., Jülicher, F., Müller, D. J. & Helenius, J. Quantification of surface tension and internal pressure generated by single mitotic cells. Sci. Rep. 4, 6213 (2014).

26. Delarue, M. et al. Compressive stress inhibits proliferation in tumor spheroids through a volume limitation. Biophys. J. 107, 1821–1828 (2014).

27. Nam, S. et al. Cell cycle progression in confining microenvironments is regulated by a growth-responsive TRPV4-PI3K/Akt-p27Kip1 signaling axis. Sci. Adv. 5, (2019).

28. Lee, H., Stowers, R. & Chaudhuri, O. Volume expansion and TRPV4 activation regulate stem cell fate in three-dimensional microenvironments. Nat. Commun. 10, 529 (2019).

29. Venkova, L. et al. A mechano-osmotic feedback couples cell volume to the rate of cell deformation. eLife 11, e72381 (2022).

30. Guo, M. et al. Cell volume change through water efflux impacts cell stiffness and stem cell fate. Proc. Natl. Acad. Sci. U. S. A. 114, E8618–E8627 (2017).

31. Okada, Y. et al. Receptor-mediated control of regulatory volume decrease (RVD) and apoptotic volume decrease (AVD). J. Physiol. 532, 3–16 (2001).

32. Jentsch, T. J. VRACs and other ion channels and transporters in the regulation of cell volume and beyond. Nat. Rev. Mol. Cell Biol. 17, 293–307 (2016).

33. Bowie, D. Shared and unique aspects of ligand- and voltage-gated ion-channel gating. J. Physiol. 596, 1829–1832 (2018).

34. Roffay, C., et al. Passive coupling of membrane tension and cell volume during active response of cells to osmosis. Proc. Natl. Acad. Sci. 118, e2103228118 (2021).

35. Marshall, W. F. Scaling of subcellular structures. Annu. Rev. Cell Dev. Biol. 36, 219–236 (2020).

36. Matthews, H. K., Bertoli, C. & de Bruin, R. A. M. Cell cycle control in cancer. Nat. Rev. Mol. Cell Biol. 23, 74–88 (2022).

37. Han, Y. L. et al. Cell swelling, softening and invasion in a three-dimensional breast cancer model. Nat. Phys. 16, 101–108 (2020).

38. Chang, J. et al. Cell volume expansion and local contractility drive collective invasion of the basement membrane in breast cancer. Nat. Mater. 23, 711–722 (2024).

39. Tsurkan, M. V. et al. Defined polymer-peptide conjugates to form cell-instructive starpeg-heparin matrices in situ. Adv. Mater. 25, 2606–2610 (2013).

40. Taubenberger, A. V. et al. 3D Microenvironment Stiffness Regulates Tumor Spheroid Growth and Mechanics via p21 and ROCK. Adv. Biosyst. 3, 1–16 (2019).

41. Mahajan, V. et al. Mapping Tumor Spheroid Mechanics in Dependence of 3D Microenvironment Stiffness and Degradability by Brillouin Microscopy. Cancers 13, 5549 (2021).

42. Welzel, P. B. et al. Macroporous StarPEG-Heparin Cryogels. Biomacromolecules 13, 2349– 2358 (2012).

43. Devany, J., Falk, M. J., Holt, L. J., Murugan, A. & Gardel, M. L. Epithelial tissue confinement inhibits cell growth and leads to volume-reducing divisions. Dev. Cell 58, 1462–1476.e8 (2023).

44. Sakaue-sawano, A. et al. Resource Visualizing Spatiotemporal Dynamics of Multicellular Cell-Cycle Progression. 487–498 (2008) doi:10.1016/j.cell.2007.12.033.

45. Panagiotakopoulou, M. et al. Cell cycle–dependent force transmission in cancer cells. Mol. Biol. Cell 29, 2528–2539 (2018).

46. Vassilev, L. T. et al. Selective small-molecule inhibitor reveals critical mitotic functions of human CDK1. Proc. Natl. Acad. Sci. 103, 10660–10665 (2006).

47. Barer, R. Interference Microscopy and Mass Determination. Nature 169, 366–367 (1952).

48. Scarcelli, G. et al. Noncontact three-dimensional mapping of intracellular hydromechanical properties by Brillouin microscopy. Nat. Methods 12, 1132–1134 (2015).

49. Schlüßler, R. et al. Correlative all-optical quantification of mass density and mechanics of sub-cellular compartments with fluorescence specificity. eLife 11, 1–23 (2022).

50. Nikolić, M., Scarcelli, G. & Tanner, K. Multimodal microscale mechanical mapping of cancer cells in complex microenvironments. Biophys. J. 121, 3586–3599 (2022).

51. Hilai, K., et al. Mechanical evolution of metastatic cancer cells in three-dimensional microenvironment. bioRxiv 1–19 (2024) doi:10.1101/2024.06.27.601015.

52. Tavares, S. et al. Actin stress fiber organization promotes cell stiffening and proliferation of pre-invasive breast cancer cells. Nat. Commun. 8, 15237 (2017).

53. Gottheil, P. et al. State of Cell Unjamming Correlates with Distant Metastasis in Cancer Patients. Phys. Rev. X 13, 031003 (2023).

54. Schmitz, A., Fischer, S. C., Mattheyer, C., Pampaloni, F. & Stelzer, E. H. K. Multiscale image analysis reveals structural heterogeneity of the cell microenvironment in homotypic spheroids. Sci. Rep. 7, 43693 (2017).

55. Ong, H. T. et al. Digitalized organoids: integrated pipeline for high-speed 3D analysis of organoid structures using multilevel segmentation and cellular topology. Nat. Methods 22, 1343–1354 (2025).

56. Alexander, M. R. et al. Regulation of Cell Cycle Progression by Swe1p and Hog1p Following Hypertonic Stress. Mol. Biol. Cell 12, 53–62 (2001).

57. Taïeb, H. M. et al. Osmotic pressure modulates single cell cycle dynamics inducing reversible growth arrest and reactivation of human metastatic cells. Sci. Rep. 11, 1–13 (2021).

58. Arsenijevic, T. et al. Hyperosmotic stress induces cell cycle arrest in retinal pigmented epithelial cells. Cell Death Dis. 4, e662 (2013).

59. Zhang, Y. et al. WNK1 is required for proliferation induced by hypotonic challenge in rat vascular smooth muscle cells. Acta Pharmacol. Sin. 39, 35–47 (2018).

60. Vahala, D. et al. 3D Volumetric Mechanosensation of MCF7 Breast Cancer Spheroids in a Linear Stiffness Gradient GelAGE. Adv. Healthc. Mater. 12, 1–13 (2023).

61. Monnier, S. et al. Effect of an osmotic stress on multicellular aggregates. Methods 94, 114–119 (2016).

62. Swaminathan, V. et al. Mechanical Stiffness Grades Metastatic Potential in Patient Tumor Cells and in Cancer Cell Lines. Cancer Res. 71, 5075–5080 (2011).

63. Alibert, C., Goud, B. & Manneville, J.-B. Are cancer cells really softer than normal cells? Biol. Cell 109, 167–189 (2017).

64. Rianna, C., Radmacher, M. & Kumar, S. Direct evidence that tumor cells soften when navigating confined spaces. Mol. Biol. Cell 31, 1726–1734 (2020).

65. Friedl, P., Wolf, K. & Lammerding, J. Nuclear mechanics during cell migration. Curr. Opin. Cell Biol. 23, 55–64 (2011).

66. Aceto, N. et al. Circulating Tumor Cell Clusters Are Oligoclonal Precursors of Breast Cancer Metastasis. Cell 158, 1110–1122 (2014).

67. Kempe, H., Schwabe, A., Crémazy, F., Verschure, P. J. & Bruggeman, F. J. The volumes and transcript counts of single cells reveal concentration homeostasis and capture biological noise. Mol. Biol. Cell 26, 797–804 (2015).

68. Padovan-Merhar, O. et al. Single Mammalian Cells Compensate for Differences in Cellular Volume and DNA Copy Number through Independent Global Transcriptional Mechanisms. Mol. Cell 58, 339–352 (2015).

69. Hertz, H. H. Hertz, Über die Berührung fester elastischer Körper, Journal für die reine und angewandte Mathematik 92, 156-171 (1881). J. Für Reine Angew. Math. 171, 156–171 (1881).

70. Sneddon, I. N. The relation between load and penetration in the axisymmetric boussinesq problem for a punch of arbitrary profile. Int. J. Eng. Sci. 3, 47–57 (1965).

71. Schlüßler, R. et al. Mechanical Mapping of Spinal Cord Growth and Repair in Living Zebrafish Larvae by Brillouin Imaging. Biophys. J. 115, 911–923 (2018).

72. Schindelin, J., et al. Fiji: An open-source platform for biological-image analysis. Nat. Methods 9, 676–682 (2012).

73. Schmidt, U., Weigert, M., Broaddus, C. & Myers, G. Cell detection with star-convex polygons. Lect. Notes Comput. Sci. Subser. Lect. Notes Artif. Intell. Lect. Notes Bioinforma. 11071 LNCS, 265–273 (2018).

74. Weigert, M., Schmidt, U., Haase, R., Sugawara, K. & Myers, G. Star-convex polyhedra for 3D object detection and segmentation in microscopy. Proc. -2020 IEEE Winter Conf. Appl. Comput. Vis. WACV 2020 3655–3662 (2020) doi:10.1109/WACV45572.2020.9093435.

75. Weigert, M. & Schmidt, U. Nuclei Instance Segmentation and Classification in Histopathology Images with Stardist. ISBIC 2022 -Int. Symp. Biomed. Imaging Chall. Proc. 2–5 (2022) doi:10.1109/ISBIC56247.2022.9854534.

76. Stringer, C., Wang, T., Michaelos, M. & Pachitariu, M. Cellpose: a generalist algorithm for cellular segmentation. Nat. Methods 18, 100–106 (2021).

77. Pachitariu, M. & Stringer, C. Cellpose 2.0: how to train your own model. Nat. Methods 19, 1634–1641 (2022).

78. Sofroniew, N., Lambert, T., Bokota, G., Nunez-Iglesias, J., Sobolewski, P., Sweet, A., Gaifas, L., Evans, K., Burt, A., Doncila Pop, D., Yamauchi, K., Weber Mendonça, M., Buckley, G., Vierdag, W.-M., Royer, L., Can Solak, A., Harrington, K. I. S., Ahlers, R. napari: a multi-dimensional image viewer for Python (v0.5.4). Zenodo (2024) doi:10.5281/zenodo.13863809.

79. von Chamier, L. et al. Democratising deep learning for microscopy with ZeroCostDL4Mic. Nat. Commun. 12, 1–18 (2021).

80. Machado, S., Mercier, V. & Chiaruttini, N. LimeSeg: A coarse-grained lipid membrane simulation for 3D image segmentation. BMC Bioinformatics 20, 1–12 (2019).

81. Cuche, E., Marquet, P. & Depeursinge, C. Spatial filtering for zero-order and twin-image elimination in digital off-axis holography. Appl. Opt. 39, 4070 (2000).

82. Wolf, E. Three-dimensional structure determination of semi-transparent objects from holographic data. Opt. Commun. 1, 153–156 (1969).

83. Sung, Y. et al. Optical diffraction tomography for high resolution live cell imaging. Opt. InfoBase Conf. Pap. 17, 1977–1979 (2009).

84. Kim, K. et al. High-resolution three-dimensional imaging of red blood cells parasitized by Plasmodium falciparum and in situ hemozoin crystals using optical diffraction tomography. J. Biomed. Opt. 19, 1 (2013).

85. Dobin, A. et al. STAR: Ultrafast universal RNA-seq aligner. Bioinformatics 29, 15–21 (2013).

86. Liao, Y., Smyth, G. K. & Shi, W. FeatureCounts: An efficient general purpose program for assigning sequence reads to genomic features. Bioinformatics 30, 923–930 (2014).

