## Supplementary data for "Cells dynamically adapt their nuclear volumes and proliferation rates during single to multicellular transitions"

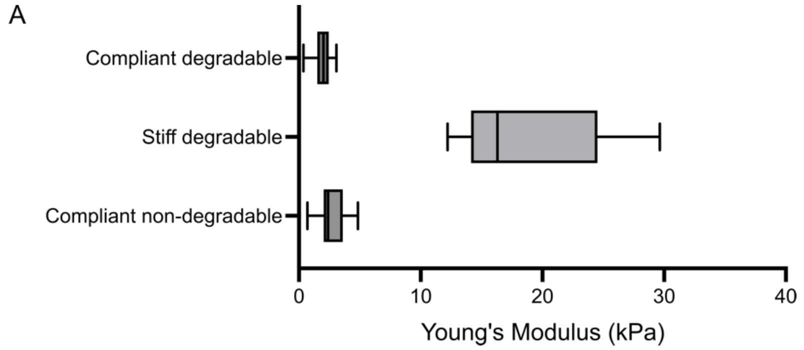

*Figure S1: (A) Stiffness (Young's modulus) of compliant degradable, stiff degradable and compliant non-degradable PEG-heparin gels as measured by AFM (atomic force microscopy). Line in the box shows the median; box shows the 25th and 75th percentile; whiskers show the minimum and maximum values. n = 20 (compliant degradable), 8 (stiff degradable), 16 (compliant non-degradable) gels.*

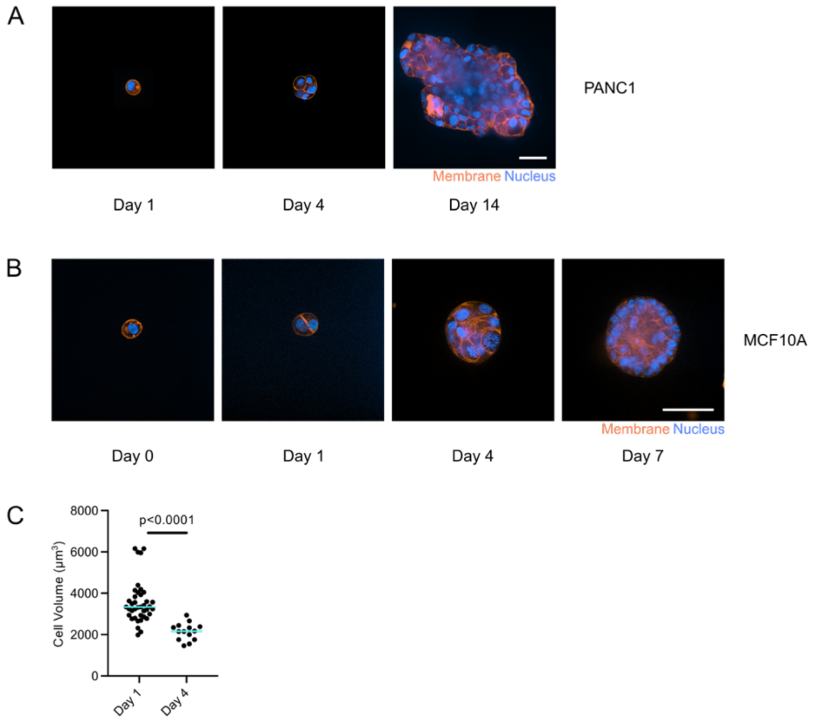

*Figure S2: (A) Live confocal images of PANC1 H2B-mCherry in compliant degradable PEG-heparin gels with CellMask Deep Red (membrane) at day 1, 4, 14. (B) Live confocal images of MCF10A expressing H2B-GFP in gels with CellMask Deep Red (membrane) at day 0, 1, 4, 7. (C) Cell volumes of HeLa FUCCI cells in gels at day 1 (absolute) and day 4 (average). n = 39 (day 1), 14 (day 4); 2-4 gels each; N = 1-2. Cyan bars represent medians. Mann-Whitney test was performed for statistical analysis. Scale bars – (A, B) 50 µm.*

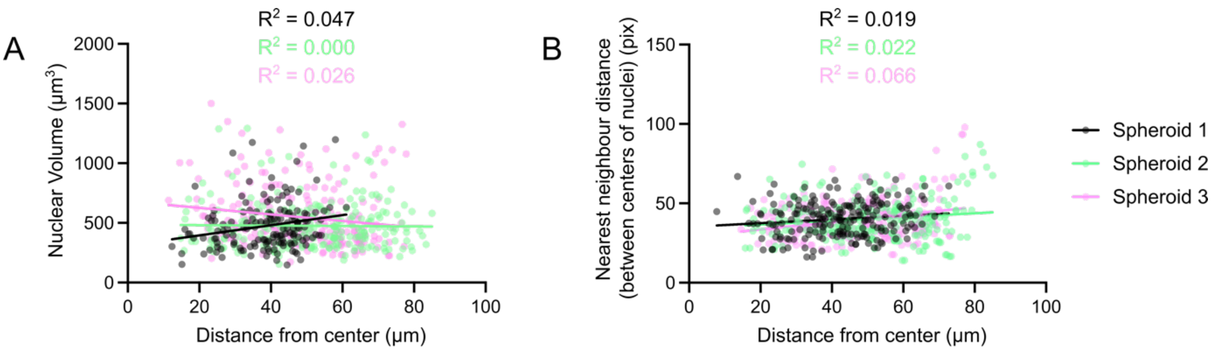

Day 14 - MCF-7 tumor spheroids

*Figure S3: (A) Nuclear volumes of MCF-7 H2B-mCherry spheroids (day 14) in compliant degradable PEG-heparin gels in relation to the distance of nuclei from the centre. n = 602; 3 spheroids. (B) Closest neighbour distance of the nuclei in MCF-7 H2B-mCherry spheroids in relation to their distance from the centre. n = 620; 3 spheroids. Linear regression was performed and goodness-of-fit was used to calculate* $R^2$.

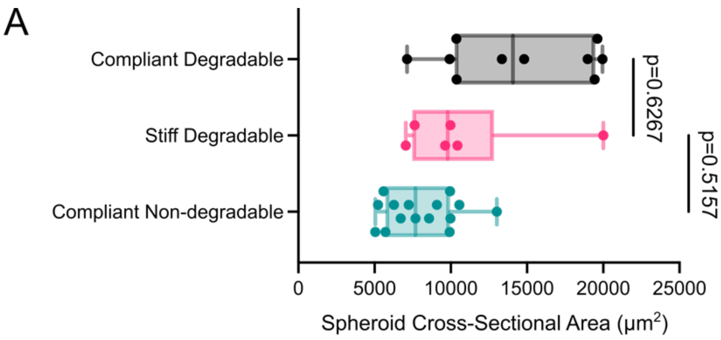

Day 14 - MCF-7 tumor spheroids

*Figure S4: (A) Cross-sectional area MCF-7 H2B-mCherry spheroids (day 14) in compliant degradable, stiff degradable and compliant non-degradable PEG-heparin gels. Line in the box shows the median; box shows the 25th and 75th percentile; whiskers show the minimum and maximum values. n = 10 (compliant degradable), 6 (stiff degradable), 15 (compliant non-degradable); 4 gels each; N = 2.*

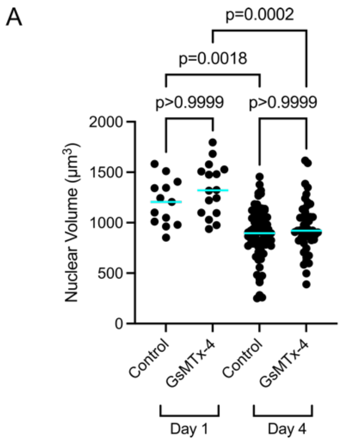

*Figure S5: (A) Nuclear volume of MCF-7 H2B-mCherry in compliant non-degradable PEG-heparin gels at day 1 and 4 under 10µm GsMTx-4 (Abcam - ab141871) (vehicle control - 41µL distilled water in 1mL media). Drug was added from the beginning. n = 13 (day 1 control), 16 (day 1 GsMTx-4), 69 (day 4*

control), 46 (day 4 GsMTx-4); 2 gels each; N = 1. Cyan bars represent the median. Kruskal-Wallis test with multiple comparisons (Dunn's) was performed for statistical analysis.

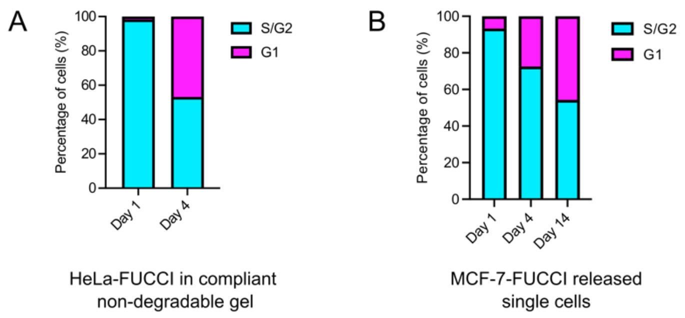

*Figure S6: (A) Percentage of HeLa FUCCI cells in G1 and S/G2 phases in compliant non-degradable PEG-heparin gels at day 1 and 4. n = 57 (day 1), 222 (day 4); 2-6 gels each; N = 1-3. (B) Percentage of MCF-7 FUCCI single cells released from compliant degradable PEG-heparin gels and multicellular structures in G1 and S/G2 at day 1, 4, 14. n = 45 (day 1), 51 (day 4), 162 (day 14); 4-6 gels each; N = 2.*

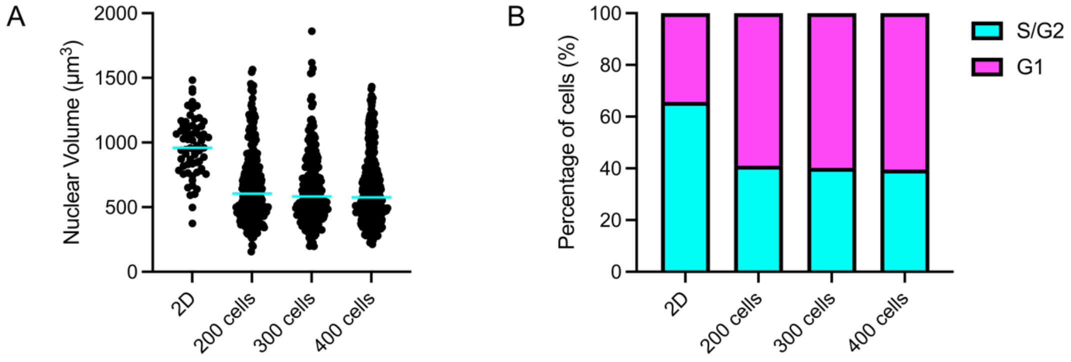

*Figure S7: (A) Nuclear volumes of MCF-7 FUCCI free-floating aggregates (200, 300, 400 cells) and single cells taken from 2D culture. n = 63 (2D), 301 (200 cells), 238 (300 cells), 378 (400 cells); 4 aggregates each; N=1. (B) Percentage of MCF-7 FUCCI cells in G1 and S/G2 phases in free-floating aggregates composed of 200, 300, 400 cells. n = 67 (2D), 352 (200 cells), 336 (300 cells), 504 (400 cells); 4 aggregates each; N=1.*

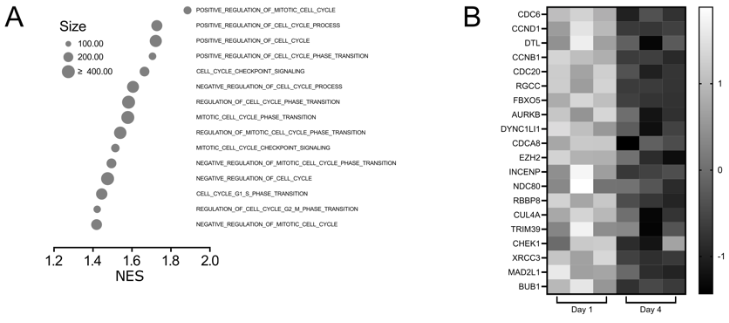

*Figure S8: (A) Gene sets related to cell cycle obtained through GSEA with NOM p-val < 0.05 and FDR q-val < 0.25. (B) Heatmap of genes involved in cell cycle (obtained from leading edge analysis of cell cycle related gene sets) which are differentially expressed between MCF-7 at day 1 and day 4 in compliant degradable PEG-heparin gels.*

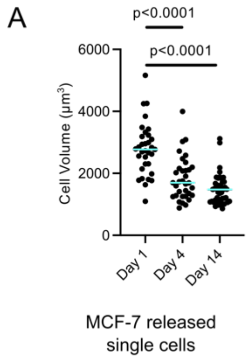

*Figure S9: (A) Cell volumes of released single MCF-7 H2B-mCherry from compliant degradable PEG-heparin gels at different culture timepoints. n = 33 (day 1), 34 (day 4), 45 (day 14); 4-6 gels each; N = 2. Cyan bars represent the median. Kruskal-Wallis test with multiple comparisons (Dunn's) was performed for statistical analysis.*

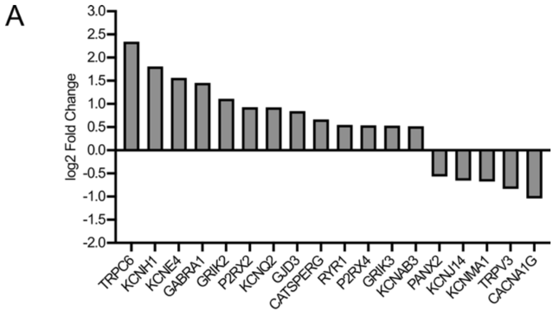

*Figure S10: (A) Fold change of genes for cellular pumps/channels being differentially expressed between MCF-7 at day 1 and 4 in compliant degradable PEG-heparin gels.*

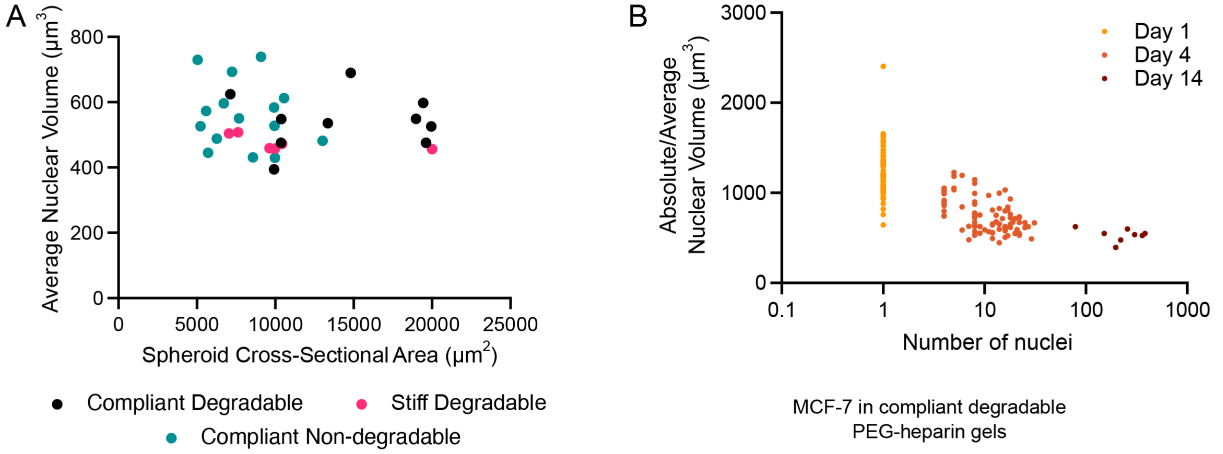

*Figure S11: (A) Average nuclear volume in relation to spheroid area for MCF-7 H2B-mCherry spheroids (day 14) in compliant and stiff degradable and compliant non-degradable PEG-heparin gels. n = 10 (compliant degradable), 6 (stiff degradable), 15 (compliant non-degradable); 4 gels each; N = 2. (B) Average (absolute for single cells) nuclear volume in relation to number of cells for MCF-7 H2B-mCherry (day 1, 4, 14) in compliant degradable PEG-heparin gels. n = 53 (day 1), 89 (day 4), 8 (day 14); 4-10 gels each; N = 2-5. Linear regression was performed and goodness-of-fit was used to calculate $R^2$.*

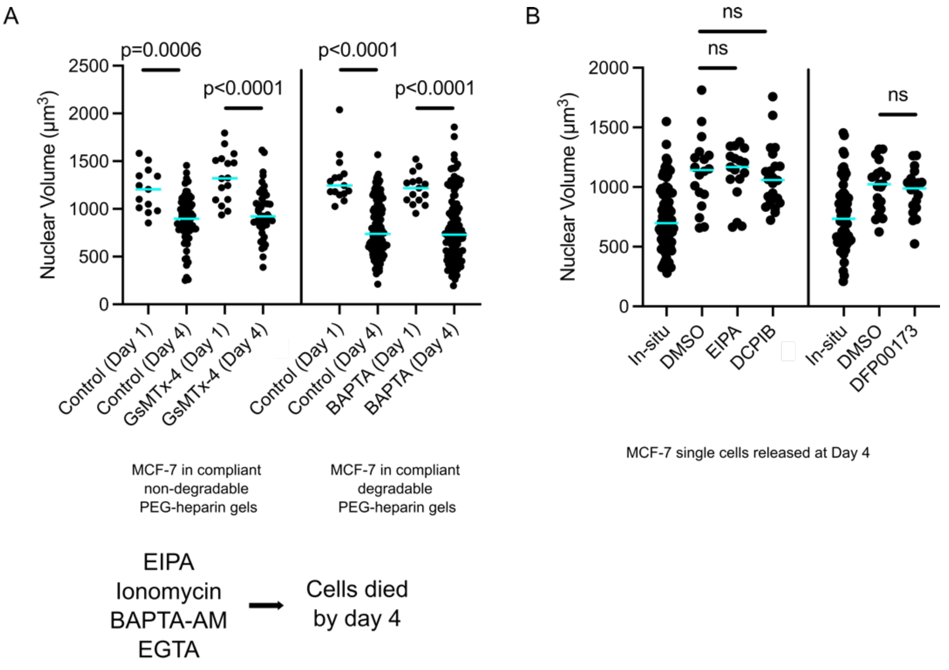

*Figure S12: (A) Nuclear volumes of MCF-7 H2B-mCherry cultured with 10 μm GsMTX-4 and 10 μm BAPTA from day 0 in compliant non-degradable and degradable (respectively) PEG-heparin gels at day 1 and 4. n=13-16 cells for day 1 and n=46-131 cells for day 4; 2 gels each; N = 1. Cells treated with 10 μm EIPA, 10 μm BAPTA-AM, 2 μm ionomycin and 5 mM EGTA did not survive till day 4. (B) Nuclear volumes of MCF-7 H2B-mCherry single cells released from compliant degradable PEG-heparin gels at day 4 along with 10 μm EIPA, 25 μm DCPIB and 10 μm DFP00173. n=17-22 cells for released single cells and 66-90 cells for in-situ control; 2 gels each; N = 1. Cyan bars represent the median. Kruskal-Wallis test with multiple comparisons (Dunn's) was performed for statistical analysis.*

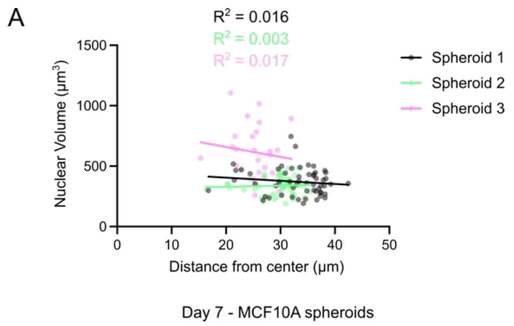

*Figure S13: (A) Nuclear volume of MCF10A H2B-GFP spheroids (day 7) in compliant degradable PEG-heparin gels in relation to the distance of nuclei from the centre. n = 139; 3 spheroids. Linear regression was performed and goodness-of-fit was used to calculate $R^2$.*

*Table 1: Cell cycle phase (G1 and S/G2) lengths obtained from light sheet imaging of MCF-7 FUCCI in compliant degradable PEG-heparin gels during days 1-4 and days 8-10 of culture time.*

| Day 2 to 4 | G1 length (hours) | | S/G2 length (hours) | Day 8 to 10 | G1 length (hours) | | S/G2 length (hours) |
|---|---|---|---|---|---|---|---|
| Entity 1 | | Entity 1 | | Entity 1 | | Entity 1 | |
| Cell 1 | 9 | Cell 1 | 18.5 | Cell 1 | 23 | Cell 1 | >17 |
| Cell 2 | 6.5 | Cell 2 | 17.5 | Cell 2 | 26.5 | Cell 2 | >23.5 |
| Cell 3 | 7.5 | Cell 3 | 22 | Cell 3 | >42.5 | Cell 3 | >22.5 |
| Cell 4 | 7.5 | | | Cell 4 | >22 | Cell 4 | 20.5 |
| Cell 5 | 9 | Entity 2 | | | | | |
| Cell 6 | >19 | Cell 1 | 15.5 | Entity 2 | | Entity 2 | |
| | | Cell 2 | 16 | Cell 1 | >14 | Cell 1 | 23 |
| Entity 2 | | Cell 3 | 19 | Cell 2 | >17 | Cell 2 | 27 |
| Cell 1 | >45.5 | | | Cell 3 | >48 | Cell 3 | 22 |
| | | Entity 3 | | | | | |
| Entity 3 | | Cell 1 | >13.5 | | | | |
| Cell 1 | 10 | Cell 2 | 24 | | | | |
| Cell 2 | 10 | | | | | | |
| Cell 3 | 9 | | | | | | |
| Cell 4 | 7 | | | | | | |
| Entity 4 | | | | | | | |
| Cell 1 | 12.5 | | | | | | |
| Cell 2 | 12.5 | | | | | | |
| Cell 3 | >32.5 | | | | | | |
